# Potent Bacterial Vaccines Require FcγRIIB-mediated Pathogen Capture by Liver Sinusoidal Endothelium

**DOI:** 10.1101/2022.07.19.500551

**Authors:** Juanjuan Wang, Haoran An, Ming Ding, Yanhong Liu, Shaomeng Wang, Qian Jin, Haodi Dong, Xianbin Tian, Jiankai Liu, Jingfei Zhang, Tao Zhu, Junqiang Li, Zhujun Shao, David E. Briles, Haifa Zheng, Linqi Zhang, Jing-Ren Zhang

**Affiliations:** Center for Infectious Disease Research, Department of Basic Medical Science, School of Medicine, Tsinghua University, Beijing, China; Tsinghua-Peking Center for Life Sciences, Tsinghua University, Beijing, China; Minhai Biotechnology, Beijing, China; Cansino Biologics, Tianjin, China; National Institute for Communicable Disease Control and Prevention, Chinese Center for Disease Control and Prevention, Beijing, China; Department of Microbiology, University of Alabama at Birmingham, Birmingham, Alabama, USA

**Author notes:** Corresponding author: Email: Haifa Zheng, Linqi Zhang or Jing-Ren Zhang. These authors contributed equally to this work (Juanjuan Wang, Haoran An, Ming Ding).

**Keywords:** Vaccine, pathogen clearance, liver, Kupffer cell, IgM, complement system, CRIg, liver sinusoidal endothelial cell, IgG, FcγRIIB, and pathogen capture

## Abstract

Certain vaccines are more effective than others against microbial infections, but the molecular mechanisms separating the two types of vaccines are largely undefined. Here, by comparing two vaccines of *Streptococcus pneumoniae* with identical antigens but different efficacies (pneumococcal conjugate vaccine – PCV13 and pneumococcal polysaccharide vaccine – PPV23), we reveal that superior vaccine protection against blood-borne bacteria is primarily achieved by activating pathogen capture of the sinusoidal endothelial cells (ECs) in the liver. Consistent with its superior protection in humans, PCV13 confers a more potent protection than PPV23 against pneumococcal infection in mice. *In vivo* real-time imaging and genetic mutagenesis revealed that PCV13 activates both liver ECs and resident macrophages Kupffer cells (KCs) to capture IgG-coated bacteria via IgG Fc gamma receptor (FcγR). In particular, the FcγRIIB-mediated capture by ECs is responsible for PCV13-induced superior protection. In contrast, PPV23 only activates KCs (but not ECs) to achieve a less effective pathogen capture and protection through complement receptor-mediated recognition of IgM- and C3-coated bacteria. These liver-based vaccine protection mechanisms are also found with the vaccines of *Neisseria meningitidis* and *Klebsiella pneumoniae*, another two important invasive human pathogens. Our findings have uncovered a novel EC- and FcγRIIB-mediated mechanism in the liver for more efficacious vaccine protection. These findings can serve as *in vivo* functional readouts to evaluate vaccine efficacy and guide the future vaccine development.

**One Sentence Summary:** Vaccine efficacy is defined by FcγRIIB-mediated capture of antibody-coated bacteria via liver sinusoidal endothelial cells.

## INTRODUCTION

Infectious diseases are a major cause of morbidity and mortality worldwide. The global landscape of infectious diseases has been continuously shaped by protection efficacy of vaccines against various pathogens. A few diseases have been globally eradicated (e.g. smallpox) or nearly eradicated (e.g. polio) by vaccination, but many other diseases are still not preventable by vaccination after many unsuccessful efforts have been made in developing effective vaccines for virtually all of the important diseases, such as malaria, tuberculosis and acquired immunodeficiency virus (AIDS) (*1*). Protection efficacy typically depends on two functional aspects of vaccine immunity: 1) the level of vaccine-induced antibody and/or T-cell responses, and 2) the potency of vaccine-elicited pathogen elimination against infection (*1*). The current knowledge on vaccine efficacy is mainly concentrated on the induction of antibody and/or T-cell responses, such as antigen selection and formulation, adjuvant effect and vaccine delivery (*2*). By contrast, it is largely unclear how vaccine-elicited immune responses contribute to pathogen clearance in the process of infection, particularly for vaccine-activated protection against blood-borne bacterial infections, the deadliest form of infections (*3*). It is believed that antibody-mediated phagocytosis by neutrophils and other circulating immune cells accounts for the elimination of blood-borne virulent bacteria in vaccinated individuals (*4, 5*), which is reflected by the use of *in vitro* phagocytosis of polymorphonuclear leukocytes for evaluation of vaccine immunity against invasive bacteria, such as *Streptococcus pneumoniae* (pneumococcus) (*6, 7*) and *Streptococcus agalactiae* (*8*).

The liver is long recognized as a major organ to clear blood-borne microbes (*9–14*). The liver resident macrophage Kupffer cells (KCs) and sinusoidal endothelial cells (ECs), representing approximately 20% and 50% of the non-parenchymal cells (NPCs), respectively, are the primary scavengers (*15*). KCs are specialized for the clearance of large particles (>0.5 µm) by actin-powered phagocytosis (*16*), whereas ECs are known to take up small particles (<0.3 µm) by clathrin-mediated endocytosis (*17*). KCs represent approximately 90% of total resident macrophages in the body (*18*). The previous studies have demonstrated that KCs are solely responsible for capturing blood-borne bacteria (*19–26*), fungi (*27*) and parasites (*28*). Our recent study reveals that certain high-virulence bacterial pathogens are able to bypass KC’s surveillance in the liver and thereby cause severe bacteremia and death (*29*). Liver ECs are regarded as a sink for viruses in the blood due to their superior capacity to KCs in the clearance of viruses (*30–33*) and bacteriophages (*34*). However, there is no evidence that ECs capture large particles with the sizes of bacteria (>1 µm), which supports the paradigm that ECs are specialized for the clearance of small particles (*17*). The earlier studies have shown that antibody- and complement-opsonized bacteria are trapped in the liver from the bloodstream (*13, 35*). However, it is unknown if the liver or its scavenging cells (KCs and ECs) plays any role in vaccine protection efficacy.

*S. pneumoniae*, an encapsulated bacterium, is a major cause of pneumonia, otitis media, bacteremia and meningitis in human. Two forms of capsular polysaccharide (CPS)-based vaccines are globally licensed to prevent invasive pneumococcal diseases (IPD) (e.g. bacteremia and meningitis): pneumococcal polysaccharide vaccine (PPV) and pneumococcal conjugate vaccine (PCV) (*4*). PPV, the first generation of pneumococcal vaccine, is composed of T cell-independent CPS antigens that primarily induce IgM antibodies, with poor immunogenicity in children under two years of ages (*36*). PCV was later developed for young children by chemical conjugation of CPSs to carrier proteins (e.g. diphtheria and anthrax toxoids), which activates both B and T lymphocytes to produce IgG antibodies (*37*). Although the current 23-valent PPV (PPV23) and 13-valent PCV (PCV13) share the same CPSs of 12 serotypes, PCV13 is shown to induce greater *in vitro* opsonophagocytosis than PPV23 (*7*), and confers a superior protection against IPD (*4*). However, except for the characteristics of vaccine-induced antibodies, the molecular mechanisms behind the differences in immunoprotection of the two vaccines remain to be defined.

To understand how effective vaccines achieve superior function in eliminating pathogens in the process of infection, this work has comprehensively delineated the vaccine-activated cellular and molecular actions in response to invasive infections by taking advantage of PCV13 and PPV23 (with common antigens but different protection efficacy) (*4*). Our data uncovered the liver as the major organ to execute vaccine-activated pathogen capture at the very early stage of bacterial blood-borne infections in mice. At the cellular level, our *in vivo* real-time imaging revealed that PCV13 and PPV23 each activates a distinct cellular mechanism of pathogen capture in the liver sinusoids, which explains their striking differences in protection efficacy. We further identified key immune molecules that are differentially engaged by PCV13 and PPV23 for pathogen capture and killing in the liver. We finally verified the two vaccine-activated mechanisms of pathogen capture in the liver with *Neisseria meningitidis* and *Klebsiella pneumoniae* vaccines. Potential application of our findings as protection markers in future vaccine development is discussed.

## RESULTS

### PCV13 is superior to PPV23 in protection against lethal pneumococcal infection

To understand how effective vaccines achieve pathogen clearance in the process of infection, we firstly assessed the immunoprotection efficacy of PCV13 and PPV23 in septic infection mouse model (**Fig. 1A**). Pneumococcal vaccines PCV13 and PPV23 have 12 identical CPS antigens but demonstrated substantially different efficacy in humans (*4*). As compared with the full mortality in the naïve group, PCV13-immunized mice survived intraperitoneal (i.p.) challenge with 10^6^ colony forming units (CFU) of serotype-4 TIGR4 pneumococci (**Fig. 1B,** left panel), a virulent strain with a median lethal dose (LD_50_) of 10^2^ CFU (*38*). Further trials revealed the full protection of PCV13 against infection of 10^7^ CFU and 2 × 10^7^ CFU, but not 10^8^ CFU of the pneumococci, yielding an LD_50_ of 4.9 ×10^7^ CFU. This result showed that PCV13 immunization is able to move up the LD_50_ value of TIGR4 by approximately 490,000 folds. In keeping with the survival rates, the protected animals displayed minimal or no bacteremia (**Fig. 1B,** right panel). By comparison, PPV23-immunized mice fully survived the infection with 10^4^ CFU of TIGR4, but further increase of infection dose led to partial (10^5^ CFU) or completely loss (10^6^ CFU) of protection (**Fig. 1C,** left panel), with an LD_50_ of 1.1 × 10^5^ CFU. The immunoprotection states against various doses of the pathogen was mirrored by dose-dependent differences in bacteremia in the first 12 hr post infection (**Fig. 1C,** right panel). Together, these results reveal that PCV13 confers 445-fold higher protection against lethal infection of pneumococci than PPV23.

**Fig. 1.**
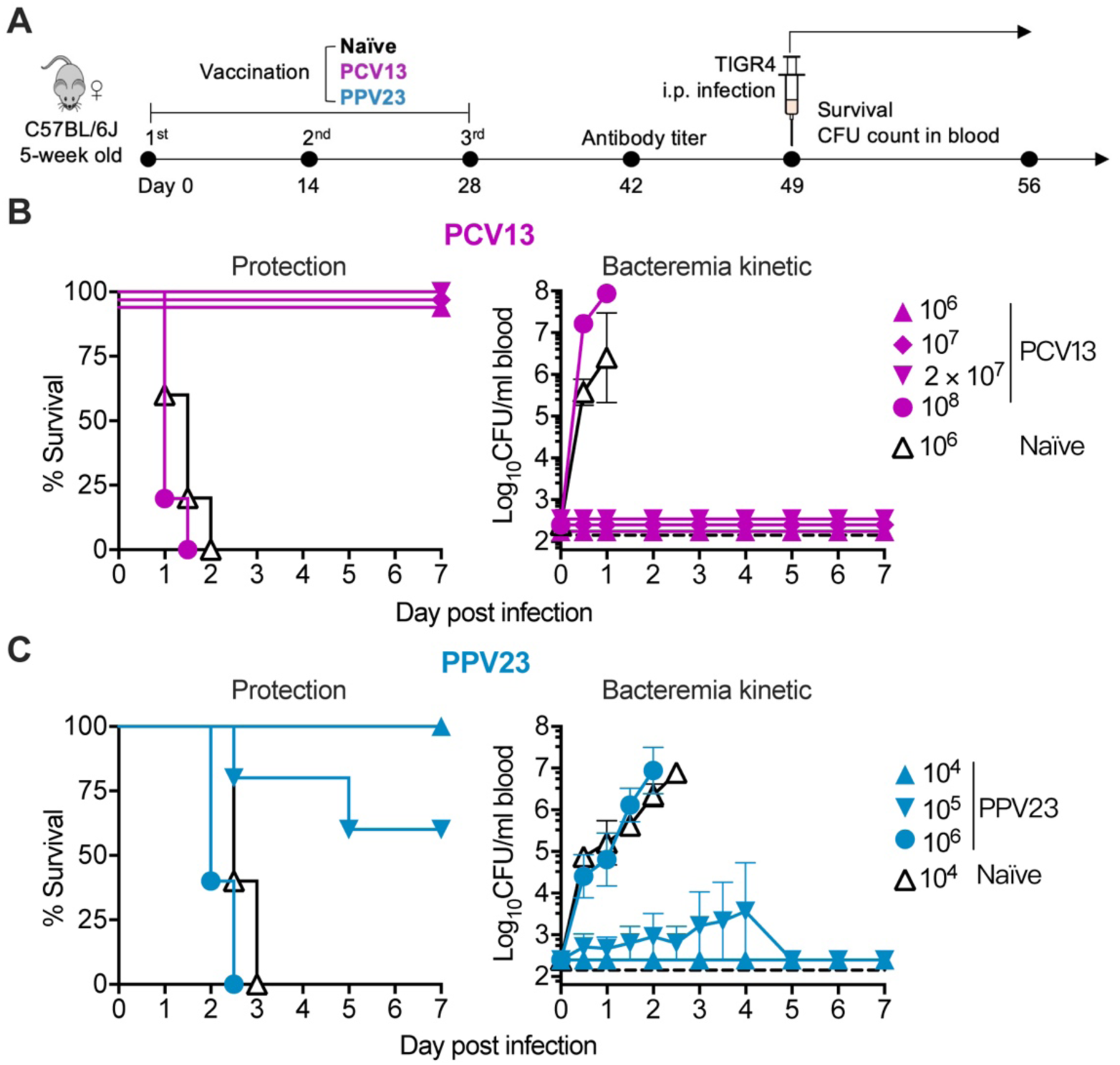
Superior protection of PCV13 over PPV23 against lethal pneumococcal infection. (**A**) Experimental design. Mice were remained untreated (naïve), or immunized with vaccine (PCV13 or PPV23) on day 0, followed by two consecutive boosts on day 14 and 28, tested for serum antibodies response on day 42, intraperitoneally (i.p.) infected with various colony-forming unit (CFU) of serotype-4 pneumococci TIGR4 on day 49, and monitored for bacteria in the blood (bacteremia kinetic) and survival for 7 days. n = 5-6. (**B** and **C**) Survival (left panel) and bacteremia kinetic (right panel) of PCV13-(B) or PPV23-immunized (C) mice were compared with naïve mice after infection. Data presented as the mean values ± standard error of mean (SEM); dotted line indicates the limit of detection.

### Superior protection is associated with faster capture and killing of pneumococci in liver

Based on our recent finding that effective pathogen clearance in the early phase of blood infections is essential for host survival (*29*), we compared the ability of PCV13 and PPV23 in clearing blood-borne pneumococci. After i.v. infection with 10^6^ CFU of TIGR4, all of naïve mice succumbed to the infection, but PCV13- and PPV23-immunized mice survived (**fig. S1A**). In contrast to sustained bacteremia in naïve mice, PCV13- and PPV23-immunized mice effectively cleared the pneumococci from the circulation in the first 30 minutes (min) post infection (**Fig. 2A**). However, PCV13-immunized mice showed a slightly more potent clearance of the pathogen than the PPV23-immunized mice, which was manifested by a 61.3-fold more bacteria in the blood of PPV23-immunized mice at 30 min (**Fig. 2A**; **fig. S1B**). This difference was also corroborated by a shorter CT_50_ value for PCV13 group (1.0 min), when 50% inoculum was removed from the blood, than the PPV23 counterpart (3.2 min) (**Fig. 2A**; **fig. S1C**). These data demonstrated that PCV13 induced faster clearance of blood-borne pathogen than PPV23.

**Fig. 2.**
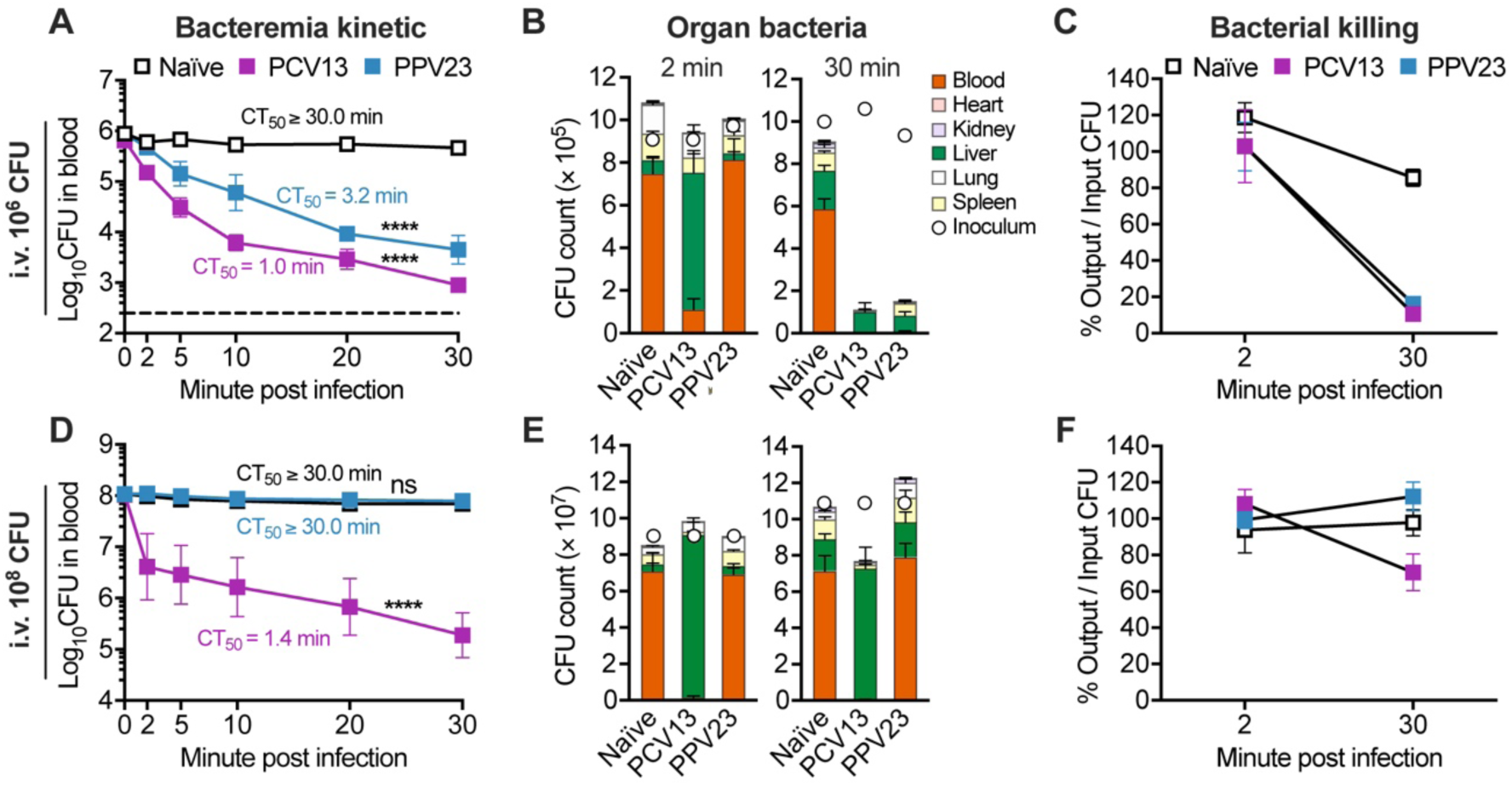
Faster pathogen capture and killing in livers of PCV13-immunized mice. (**A-C**) Vaccine-activated clearance of low-dose serotype-4 pneumococci. Naïve and PCV13- or PPV23-immunized C57BL/6J mice were tested for bacterial CFU counts in the blood (bacteremia kinetic) in the first 30 min post intravenously (i.v.) infection with 10^6^ CFU of TIGR4 (A). The time for the clearance of 50% bacteria in the blood (CT_50_) is presented for each group. CFU counts in the blood and major organs of each animal (organ bacteria) were determined at 2 or 30 min (B). Total bacteria detected in all the six samples are presented as the percentage of the inoculum at 2 and 30 min (C), which reflects *in vivo* bacterial killing during the first 30 min post infection. n = 3. (**D**-**F**) Vaccine-activated clearance of high-dose serotype-4 pneumococci. Mice were immunized and subsequently infected with 10^8^ CFU of TIGR4 to determine bacteremia kinetic (D), organ bacteria (E), and bacterial killing (F) as in (A-C). n = 3. Data presented as the mean ± SEM. Dotted line indicates the limit of detection. Statistics: (A and D) two-way ANOVA.

While vaccines are known to enhance bacterial clearance in the blood, it is believed that blood-borne pathogens are eliminated by circulating phagocytes (*4, 5*). To determine where inoculated pneumococci were captured, we measured bacterial burden in the heart, kidney, liver, lung and spleen at 2 and 30 min post infection. Consistent with rapid reduction in bacteremia (**Fig. 2A**), PCV13-immunized mice captured about 69.4% of the inoculum in the liver at 2 min post infection (**Fig. 2B**, left panel). By contrast, PPV23-immunized, like those naïve mice, showed minimal pneumococci capture in all organs tested including the liver. At 30 min post infection, however, PPV23-immunized mice appeared to improve capture efficiency in the liver and spleen, but remained significant difference from that of PCV13-immunized mice (**Fig. 2B**, right panel). For example, PCV13-immunized mice captured majority of the pneumococci in the liver (89.0%), whereas PPV23-immunized mice in both the liver (50.5%) and spleen (39.6%). These data demonstrate that PCV13 is superior to PPV23 in driving hepatic capture of pneumococci.

To determine if vaccine-activated pathogen capture leads to bacterial killing, we analyzed the total viable bacteria in the blood and five major organs of vaccinated mice. Although the majority of the 10^6^ CFU inoculum was already sequestered in the liver of PCV13-immunized mice at 2 min (**Fig. 2B**), these animals still carried a similar level of viable bacteria as the inoculum, indicating that bacteria captured by the liver were not killed immediately (**Fig. 2C**). However, the most of the inoculum became undetectable in the PCV13-immunized mice (by 89.3%) at 30 min, as compared with a marginal reduction of bacterial burden in naïve mice (by 9.3%) (**Fig. 2C**; **fig. S1D**). To a slightly less extent, the bacteria inoculated in PPV23-immunized mice were also eliminated in the first 30 min post infection (by 83.8%). Furthermore, similar findings against serotype-3 pneumococci ST865, another serotype covered by both the vaccines, were also noticed in terms of faster bacterial clearance from bloodstream (**fig. S1E**), more potent hepatic bacterial capture (**fig. S1F**), and more significant bacterial killing in PCV13-than PPV23-immunized mice (**fig. S1G-H**). These results imply that PCV13 and PPV23 accomplish the protective immunity by activating pathogen capture and killing in the liver.

As PCV13 displayed a much higher level of protection than PPV23 (**Fig. 1**), we used a higher dose (10^8^ CFU) of TIGR4 to determine if this functional difference was caused by higher capacity of pathogen clearance in the livers of PCV13-immunized mice. While PCV13-immunized mice effectively cleared the high inoculum of the pathogen at 30 min with a CT_50_ of 1.4 min, the PPV23 immunization failed to yield obvious impact on the clearance of 10^8^ CFU inoculum (**Fig. 2D**; **fig. S1I-J**). PPV23-immunized and naïve mice exhibited a similar pattern of severe bacteremia up to the first 30 min post infection (**Fig. 2D**; **fig. S1I-J**). Enumeration of organ-specific bacteria revealed highly effective capture of the pathogen in the livers of PCV13-immunized mice at 2 min (**Fig. 2E**, left panel), and maintained up to 30 min (**Fig. 2E**, right panel). To a less extent, the total variable bacteria in the PCV13 group were reduced by 29.4% at 30 min (**Fig. 2F**). By comparison, PPV23-immunized mice displayed poor pneumococci capture as naïve mice at both 2 min and 30 min, with the majority of 10^8^ CFU pneumococci in the bloodstream throughout the first 30 min. Accordingly, no significant killing of the high inoculum was observed in PPV23-immunized and naïve mice in this period (**Fig. 2F**; **fig. S1K**). Taken together, these data demonstrate that the adaptive immunity activated by PCV13 in the liver is far more potent in capturing and killing virulent pneumococci in the blood circulation.

### Faster pneumococcal capture is mainly achieved through liver sinusoidal endothelial cells

To identify hepatic cells responsible for vaccine-activated pneumococcal capture in the liver, we selectively depleted macrophages, neutrophils and monocytes using the combination of clodronate liposomes (CLL, targeting macrophage) and Gr1 antibody (targeting neutrophils and monocytes) (*39*) (**Fig. 3A**). The depletion efficacy of phagocytes was confirmed by flow cytometry and *Staphylococcus aureus* infection data obtained with the CLL/Gr1-treated mice (**fig. S2**). To our surprise, CLL/Gr1 treatment had no detectable impact on the PCV13-elicited protection. All of the depleted PCV13-immunized mice survived the lethal infection of serotype-4 pneumococci as those animals without depletion (**Fig. 3B**; **fig. S3A**). In keeping with this observation, phagocytes depletion in PCV13-immunized mice did not yield significant impact on the early clearance (**Fig. 3C**), hepatic capture (**Fig. 3D**) or killing (**Fig. 3E**) of the pneumococci. These results indicate that the dispensability of major phagocytes (macrophages, neutrophils and monocytes) in the PCV13-elicited immunity.

**Fig. 3.**
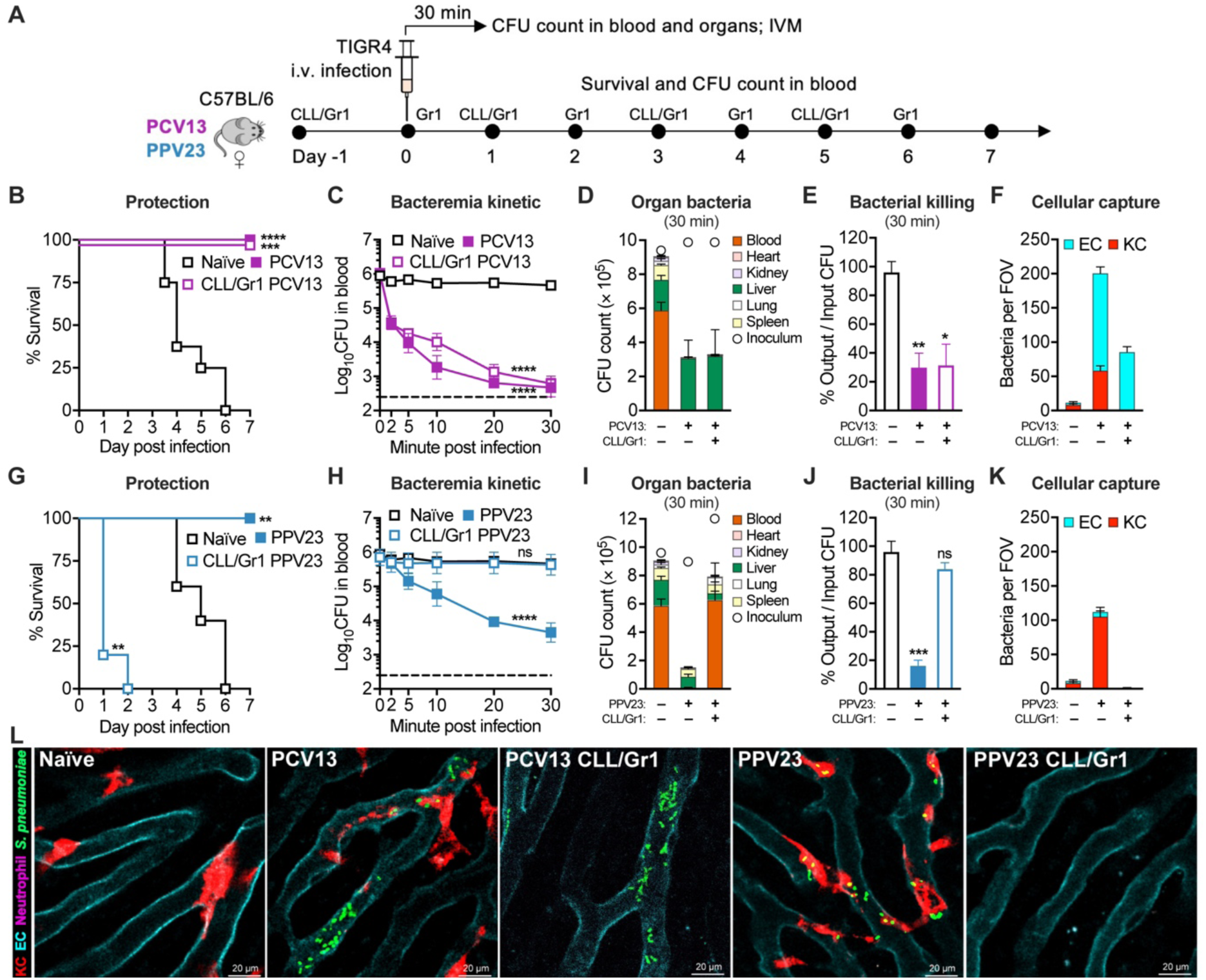
PCV13- and PPV23-specific modes of pathogen capture and killing in the liver. (**A**) Experimental design. Phagocytes were periodically depleted in PCV13- or PPV23-immunized mice by clodronate liposomes (CLL, targeting macrophage) and Gr1 antibody (targeting neutrophils and monocytes) before and after infection to assess the role of phagocytes in vaccine-activated immunity. (**B**) Dispensability of phagocytes in PCV13-elicited immunoprotection. PCV13-immunized mice were treated with CLL and Gr1 (PCV13 CLL/Gr1) and compared with untreated counterpart (PCV13) and naïve mice in survival post infection with 10^6^ CFU of TIGR4. n = 7-8. (**C-E**) Dispensability of phagocytes in PCV13-elicited pathogen clearance. PCV13-immunized mice were treated with CLL and Gr1 (PCV13 CLL/Gr1) and compared with untreated counterpart (PCV13) and naïve mice in the early clearance (C), hepatic capture (D), and killing (E) of the pathogen in the first 30 min post infection with 10^6^ CFU of TIGR4. n = 3. (**F**) The *in vivo* real-time process of PCV13-activated pathogen capture in the liver. Bacterial traffic and capture in the liver sinusoids were visualized by intravital microscopy (IVM) in PCV13-immunized mice (PCV13), the CLL/Gr1-treated counterpart (PCV13 CLL/Gr1) and naïve mice 10-15 min after i.v. infection with 5 × 10^7^ CFU of TIGR4. Bacteria associated with sinusoidal endothelial cells (ECs) and Kupffer cells (KCs) in representative images as shown in (L) are quantified and presented as the mean of bacteria in at least five random field of views (FOVs). The process of bacterial capture is demonstrated in **Movie S1** and **S2**. Quantitation was shown as pooled data of 5-16 FOVs from two mice in each group. (**G**) Essentiality of phagocytes in PPV23-elicited immunoprotection. PPV23-immunized mice were treated with CLL/Gr1 (PPV23 CLL/Gr1) and compared with untreated counterpart (PPV23) and naïve mice in survival post infection with 10^6^ CFU of TIGR4. n = 5. (**H**-**J**) Importance of phagocytes in the PPV23-activated pathogen clearance. PPV23-immunized mice (PPV23) were compared with CLL/Gr1-treated counterpart (PPV23 CLL/Gr1) and naïve mice in early clearance (H), hepatic capture (I), and killing (J) of the pathogen as in (C-E). n = 3. (**K**) The *in vivo* real-time process of PPV23-activated pathogen capture in the liver. EC- and KC-associated bacteria were quantified by IVM imaging in vaccinated mice lacking major phagocytes as in (F), except for using the PPV23 vaccine. The process of bacterial capture is demonstrated in **Movie S1** and **S2**. Quantitation was shown as pooled data of 5-10 FOVs from two mice in each group. (**L**) Representative images of vaccine-activated pathogen capture in the liver sinusoids. Pneumococci (green), KCs (red), ECs (cyan), and neutrophils (magenta) were visualized by IVM in naïve and immunized mice without or with (CLL/Gr1) phagocytes depletion. Data are presented as the mean ± SEM; dotted line indicates the limit of detection. Statistics: (B and G) log-rank (Mantel-Cox) test; (C and H) two-way ANOVA; (E and J) unpaired two-tailed Student’s t test.

To further identify the pathogen-capturing cells in the livers of PCV13-immunized mice, we visualized the behavior of circulating pneumococci in the liver sinusoids by time-lapse intravital microscopy (IVM) imaging. In the first 20 min upon i.v. inoculation, serotype-4 pneumococci smoothly trafficked through the vasculatures of naïve mice (**Fig. 3F** and **3L**; **Movie S1**), a typical feature of high-virulence pneumococci (*29*). In sharp contrast, migrating bacteria in the liver sinusoids of PCV13-immunized mice were predominantly accumulated to liver sinusoidal endothelial cells (ECs) (71.8% of total immobilized bacteria) and, to a less extent, to liver resident macrophage Kupffer cells (KCs) (28.2%) (**Fig. 3F** and **3L; Movie S1**). Depletion of major phagocytes by CLL/Gr1 treatment had little impact on PCV13-activated bacterial attachment to ECs although the KC-mediated capture was completely ablated (**Fig. 3F** and **3L; Movie S2**). The IVM data show that PCV13-activated immunity achieves the function of pneumococcal capture in the liver predominantly by ECs and, to significantly less extent, by KCs.

In an opposite manner, phagocyte depletion made PPV23-immunized mice susceptible to an otherwise nonlethal dose of TIGR4, with 100% mortality in the CLL/Gr1-treatment group (**Fig. 3G**; **fig. S3B**). As a result, CLL/Gr1 treatment rendered the vaccinated mice incapable of shuffling bacteria from the circulation (**Fig. 3H**) to the liver (**Fig. 3I**), or killing the captured pneumococci (**Fig. 3J**). These results clearly indicated that capture and killing of pneumococci in PPV23-immunized mice were predominately mediated through major phagocytes. This notion was further supported by IVM imaging where PPV23-immunized mice showed progressive attachment of bacteria to KCs (92.8%) and occasionally to ECs (7.2%). Depletion of KCs virtually abolished pneumococcal deposition in the liver sinusoids (**Fig. 3K** and **3L**; **Movie S1** and **S2**). This vaccine type-specific immune functions were also confirmed with serotype-3 pneumococci (**fig. S3C-K**; **Movie S3**). These experiments have unequivocally demonstrated that PCV13 and PPV23 each activates a unique cellular mechanism of pathogen capture in the liver, and the PCV13-elicited faster pathogen capture is mainly executed by liver sinusoidal endothelial cells.

### Endothelial capture is determined by antibody abundance and subtypes

To understand how PCV13-induced immunity mediated capture of pneumococci in the liver, we tested whether antibody-mediated opsonophagocytosis played a role as antibody levels are correlated with protection of pneumococcal conjugate vaccines (*6, 7*). PCV13 immunization led to a high level of IgG and relatively lower level of IgM antibodies in the blood against CPS4 in WT mice. As anticipated, there was no detectable IgG or IgM in vaccinated B cell-deficient μMT mice (**Fig. 4A**). As a result, PCV13-immunized μMT mice lost the ability to shuffle pneumococci from the bloodstream (**fig. S4A**) to the liver (**Fig. 4B**; **fig. S4B**), demonstrating the essential role of antibodies in PCV13-activated pneumococcal capture in the liver. This finding was further verified with purified IgG antibodies from PCV13 immune serum, which showed a dose-dependent shuffling of antibody-opsonized pneumococci from the bloodstream to the liver of naïve mice (**fig. S4C-F**). We next studied the impact of IgG antibodies on PCV13-activated pathogen capture by ECs and KCs using PCV13 immune serum-treated pneumococci. IVM imaging revealed that the bacteria pretreated with relatively small amount of the serum (100 μl) exclusively bound to KCs without significant binding to ECs, but increase in immune serum (200-300 μl) led to a dose-dependent shift of immobilized bacteria to ECs (**Fig. 4C**; **Movie S4**). This result indicates that bacterial binding by ECs requires relatively higher abundance of antibodies than the activity of KCs.

**Fig. 4.**
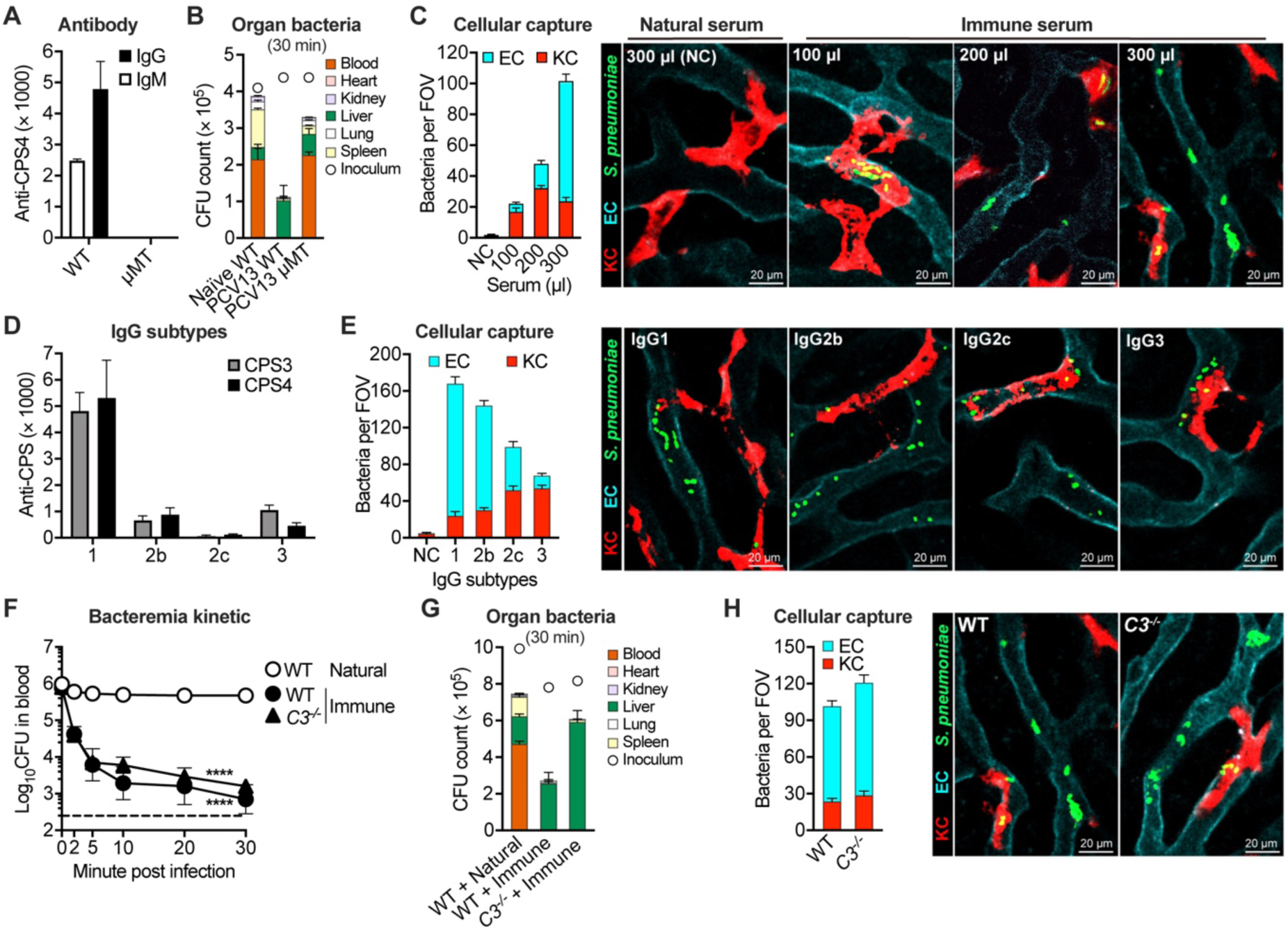
Importance of IgG abundance and subtypes in PCV13-activated pathogen capture by liver sinusoidal endothelium. (**A**) PCV13-elicited antibodies. IgG and IgM antibodies against serotype-4 CPS (anti-CPS4) were tested in sera of PCV13-immunized WT and B cell-deficient μMT mice by ELISA. n = 3-9. (**B**) Essentiality of antibodies in PCV13-elicited hepatic capture of pathogen. Bacteria in the blood and major organs were determined in PCV13-immunized WT (PCV13 WT) or the μMT counterpart (PCV13 μMT), and naïve C57BL/6J mice at 30 min post i.v. infection with 10^6^ CFU of TIGR4. n = 3. (**C**) Dose-dependent activity of PCV13 immune serum in activating pathogen capture by ECs. C57BL/6J mice were i.v. infected with 2 × 10^7^ CFU of TIGR4 that were pretreated with various amounts of heat-inactivated immune serum from PCV13-immunized mice before infection. Natural serum from naïve mice was used as a negative control (NC). Pneumococci immobilized on ECs and KCs were quantified as in Fig. 3F. The process of bacterial capture is demonstrated in **Movie S4**. Quantitation was shown as pooled data of 5 FOVs from two mice in each group. (**D**) Quantitative dominance of IgG1 antibodies elicited by PCV13 immunization. Subtypes of IgG antibodies against serotype-3 (anti-CPS3) or −4 (anti-CPS4) capsule were quantified in PCV13 immune serum as in (A). n = 3. (**E**) Functional dominance of IgG1 antibodies in activating pathogen capture by ECs. Bacteria immobilized on ECs and KCs were quantified as in Fig. 3F, except for infecting C57BL/6J mice with 5 × 10^7^ CFU of serotype-3 pneumococci ST865 before subsequent injection of purified IgG1, IgG2b, IgG2c or IgG3 variants of the 5F6 monoclonal antibody against CPS3 (2.5 mg/kg). The process of bacterial capture is demonstrated in **Movie S6**. Quantitation was shown as pooled data of 10-20 FOVs from two mice in each group. (**F-H**) Dispensability of complement C3 in PCV13 immune serum-mediated capture of TIGR4 in the liver. Bacteremia kinetic (F) and organ bacteria (G) were assessed in naïve WT and *C3*^-/-^ mice after being i.v. infected with 10^6^ CFU of 10 µl PCV13 immune serum-opsonized TIGR4; ECs- and KCs-associated bacteria (H) quantified by IVM as in (C) post i.v. infection with 300 µl PCV13 immune serum-opsonized TIGR4. Bacterial capture is demonstrated in **Movie S7**. n = 3 for (F and G); quantitation in (H) was shown as pooled data of 5-19 FOVs from two mice in each group. Data shown in (A and D) were representative of duplicate experiments. Data are presented as the mean ± SEM. Statistics: (F) two-way ANOVA.

It is known that vaccine dose (or amount) and immunization time are two key parameters for immunization schedules of all human vaccines, but immunization schedules of many vaccines are empirically based. The immunization schedule of PCV13 ranges from one to four intramuscular injections of 500 µl, depending on the age of children. We first determined the impact of vaccine dose on pathogen capture in the liver by immunizing mice once with three different doses of PCV13. This experiment showed a vaccine dose-dependent antibody production (**fig. S4G**), pathogen capture in the liver sinusoids (**fig. S4H**; **Movie S5**), and protection against lethal pneumococcal infection (**fig. S4I**). While immunization with 0.25 µl of PCV13 did not elicit any detectable anti-CPS4 antibody or pathogen capture and no protection post i.p. infection with 10^6^ CFU of TIGR4, a 10-fold increase in vaccine dose (2.5 µl) led to low levels of IgG antibodies, pathogen binding to KCs (but not ECs) and partial protection of survival (50%). Vaccination with 50 µl of PCV13, a dose regularly used for all the other immunization experiments in this work, resulted in additional increase in ant-CPS4 antibodies, immobilized bacteria on the ECs and full protection against infection. In particular, the largest vaccine dose enabled ECs to capture 62.5% of the total immobilized bacteria, indicating the importance of an adequate vaccine dose in eliciting potent pathogen capture by ECs. We next tested the impact of boost immunization using one or two boost immunizations with 2.5 µl of PCV13, a minimal dose capable of eliciting detectable antibody response and bacterial capture by KCs (**fig. S4G-I**). While vaccination once (Vac-1) yielded low serotype-specific IgG antibodies, the antibody responses were enhanced by one (Vac-2) and two (Vac-3) boost immunizations (**fig. S4J**). IVM imaging also revealed an immunization time-dependent shift of immobilized pneumococci from KCs to ECs (**fig. S4K**; **Movie S5**). While the Vac-1 mice showed KC-specific enhancement in bacterial capture, the majority of immobilized pneumococci were shifted to ECs in Vac-2 and Vac-3 mice, which confers both full protection against i.p. infection of 10^7^ CFU of TIGR4 (**fig. S4L**). These data show that an appropriate immunization schedule is essential for achieving vaccine-activated pathogen capture by ECs.

Since PCV13 predominantly elicited IgG1 with minimal levels of IgG2b and IgG3 against serotype-3 CPS (CPS3) and CPS4 (**Fig. 4D**), we chose to study the potential impact of IgG subtypes on EC’s capture of pneumococci. To this end, we synthesized IgG1, IgG2b, IgG2c and IgG3 form of 5F6, a CPS3-specific antibody able to protects mice from lethal infection of serotype-3 pneumococci (*40*), and tested their protectivity in the same model of pneumococcal infection in mice. All IgG subtypes showed a comparable *in vitro* binding to the CPS3 antigen (**fig. S5A**), and were all able to drive bacteria from the bloodstream (**fig. S5B**) to the liver (**fig. S5C**). However, IVM imaging revealed striking differences among the IgG subtypes in two related aspects: 1) the overall capture capacity of pneumococci in the liver; 2) distribution of immobilized bacteria between ECs and KCs (**Fig. 4E**; **Movie S6**). The IgG1 was outstanding in both the categories, with the highest total immobilized pneumococci per FOV and EC/KC bacterial ratio (6:1). The favoritism of IgG1-opsonized pathogen to ECs depends on the abundance of the antibody, which is reflected in the dose-dependent shift of immobilized bacteria to ECs from KCs (**fig. S5D-F**; **Movie S6**). IgG2b yielded slightly lower levels of immobilized bacteria and EC/KC bacterial ratio (4:1), but IgG2c and IgG3 variants showed significantly weaker activities in both the categories. The vast majority of IgG3-opsonized pneumococci were immobilized by KCs. This finding shows that abundant production of IgG1 antibodies and to lesser extent IgG2b are correlated with superior pneumococcal capture by ECs in PCV13-immunized mice.

Based on the importance of the complement system in antibody-mediated opsonophagocytosis of pathogens (*41*), we next studied the contribution of the system to PCV13-driven pneumococcal capture and killing using mice deficient in complement protein C3, a key component of the complement system (*41*). As previously described (*41*), PCV13 immunization in *C3*^-/-^ mice showed partially impaired IgG production (**fig. S6A**). We thus assessed the complement function in naïve mice with pneumococci opsonized by PCV13 immune serum. *C3*^-/-^ mice did not show any impairment either in shuffling antibody-opsonized pneumococci from bloodstream (**Fig. 4F**) to the liver (**Fig. 4G**), or pneumococcal capture by ECs (**Fig. 4H**; **Movie S7**). However, *C3*^-/-^ mice displayed significant deficiency in pneumococcal killing and immunoprotection (**fig. S6B-C**), which was verified with serotype-3 pneumococci (**fig. S6D-F**). The critical role of C3 in mediating killing (rather than capture) was further confirmed in PCV13-immunized *C3*^-/-^ mice (**fig. S6G-K**). These observations suggest that the complement system is dispensable for PCV13-activated pneumococcal capture by ECs, but essential for subsequent killing of EC-bound pneumococci.

### Endothelial capture is achieved through IgG1-FcγRIIB interaction

To understand how ECs recognize IgG-opsonized bacteria, we sought to identify responsible IgG Fc receptor(s). Proteomic analysis of murine ECs identified Fc**γ**RIIB and neonatal Fc receptor (FcRn) were predominantly expressed, out of the five known Fc receptors tested (**Fig. 5A**; **Table S1**). Stable expression of Fc**γ**RIIB in CHO cells (**Fig. 5B)** led to dramatic increase in adhesion of serotype-4 pneumococci after opsonized with PCV13 immune serum (**Fig. 5C**). By contrast, co-expression of FCGRT and B2m, the two subunits of FcRn, failed to enhance bacterial adhesion to CHO cells. Furthermore, opsonization by PCV13 immune serum significantly boosted pneumococcal adhesion to primary ECs only from wildtype mice but not from *Fcgr2b^-/-^* mice (**Fig. 5D**; **fig. S7**). These *in vitro* data strongly suggested that Fc**γ**RIIB but not FcRn mediates the capture of antibody-opsonized pneumococci by ECs. We next investigated relative contribution of Fc**γ**RIIB-mediated pathogen capture by ECs to the overall protection efficacy of PCV13. PCV13 immunization led to full survival of wildtype mice, but all of the vaccinated *Fcgr2b*^-/-^ mice succumbed to pneumococcal infection (**Fig. 5E,** left panel). PCV13 immunization therefore enabled the wildtype mice to successfully clear the otherwise lethal inoculum at the early stage of infection, whereas the immunized *Fcgr2b*^-/-^ mice failed to do so with severe bacteremia up to 10^7^ CFU (**Fig. 5E**, right panel). Surprisingly, the depletion of major phagocytes (macrophages, neutrophils and monocytes) had minimal impact on PCV13-boosted protection and pneumococcal clearance under this experimental condition (**Fig. 5E**). This result shows that Fc**γ**RIIB-mediated capture of IgG-opsonized pathogen by ECs contributed substantially to the superior protection efficacy of PCV13.

**Fig. 5.**
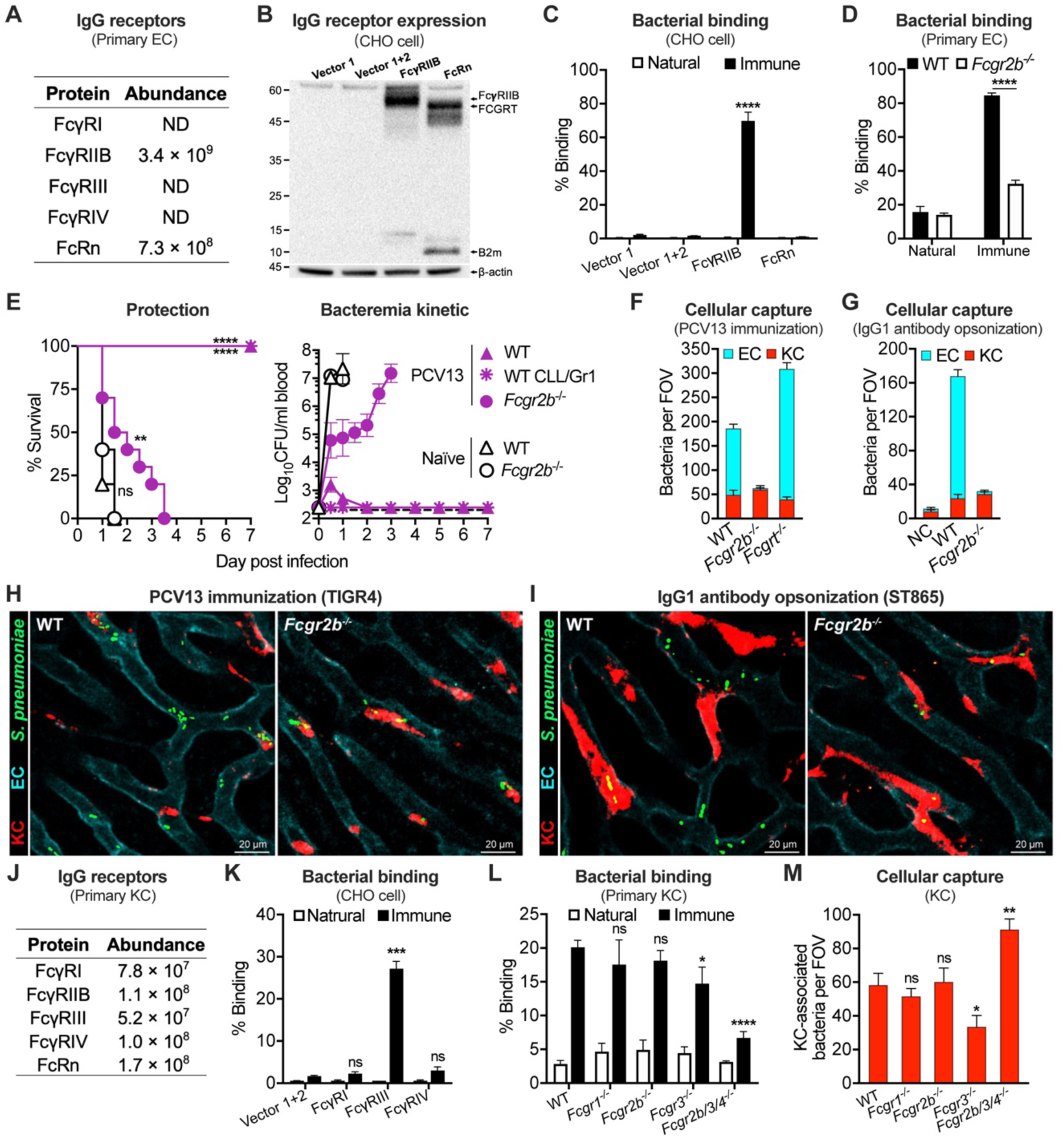
FcγRIIB-mediated capture of IgG1-coated pathogen by liver sinusoidal endothelium. (**A**) Relative abundance of IgG Fc receptors in primary murine ECs. Protein expression was detected by mass spectrometry and presented as the means of peak areas from two individual trials. ND, not detectable. Additional data are available in **Table S1**. (**B**) Stable expression of FcγRIIB (*Fcgr2b*) or FcRn (*Fcgrt* and *B2m*) in CHO cells as detected by Western blotting. The cDNA of *Fcgr2b* or *Fcgrt* were inserted in pCDH (vector 1); that of *b2m* encoding the small subunit of FcRn in pFUGW (vector 2). β actin was used as a loading control. Molecular standards are marked as kilodaltons (kDa). (**C**) *In vitro* binding of antibody-coated pneumococci to FcγRIIB-expressing CHO cells. TIGR4 was incubated with heat-inactivated natural or immune serum before cellular binding assay; the results of representative experiment presented as the percentages of cell-associated bacteria out of the total CFU at the end of the assay. n = 3. (**D**) The requirement of FcγRIIB for *in vitro* binding of antibody-coated pneumococci to primary murine ECs. The binding assay was performed and presented as in (C), except for using primary ECs of WT and *Fcgr2b*^-/-^ mice. (**E**) Superior immunoprotection potency of FcγRIIB-mediated pathogen capture by ECs. Specific contributions of PCV13-activated pathogen capture by ECs and KCs to immunoprotection are reflected in the survival (left panel) and bacteremia kinetic (right panel) of vaccinated *Fcgr2b*^-/-^ (filled circle) or WT C57BL/6J mice lacking phagocytes (asterisk) post i.p. infection with 2 × 10^7^ CFU of TIGR4, respectively. PCV13-immunized WT (filled triangle) and naïve WT (open triangle) or *Fcgr2b*^-/-^ (open circle) mice were used as controls. n = 10. (**F**) Essentiality of FcγRIIB for PCV13-activated bacterial capture by ECs. EC- and KC-associated bacteria in PCV13-immunized WT, *Fcgr2b*^-/-^ or *Fcgrt*^-/-^ C57BL/6J mice were assessed by IVM post i.v. infection with TIGR4 as in Fig. 3F. The process of bacterial capture is demonstrated in (**H**) and **Movie S8**. Quantitation was shown as pooled data of 6-15 FOVs from two mice in each group. (**G**) Requirement of FcγRIIB for IgG1-medited bacterial capture by ECs. Anti-CSP3 IgG1-opsonized bacteria in the liver sinusoids of WT and *Fcgr2b*^-/-^ C57BL/6J mice were quantified by IVM imaging post i.v. infection with ST865 as in Fig. 4E. The process of bacterial capture is demonstrated in (I) and **Movie S6**. Quantitation was shown as pooled data of 5-15 FOVs from two mice in each group. (**H** and **I**) Demonstration of FcγRIIB importance in IgG1-mediated pathogen capture by ECs. Representative IVM images of the liver sinusoids are shown to demonstrate that FcγRIIB is fully responsible for the EC’s capture of the pathogen in PCV13-immunized mice (H) or naïve mice that were treated with purified IgG1 antibody (I). Related data are also presented elsewhere in (F) and Movie S8 for the vaccinated mice, and in (G) and Movie S6 for the IgG1-treated mice. (**J**) Relative abundance of each IgG Fc receptors in murine KCs as detected by mass spectrometry as in (A). Additional data are available in **Table S2**. (**K**) *In vitro* binding of TIGR4 to CHO cells expressing IgG receptor FcγRI (*Fcgr1* and *Fcg*γ), FcγRIII (*Fcgr3* and *Fcg*γ) or FcγRIV (*Fcgr4* and *Fcg*γ) as in (C). The cDNA of *Fcgr1, 3 or 4* was inserted in pCDH (vector 1); that of *Fcg*γ encoding the common γ chain in pFUGW (vector 2). (**L** and **M**) Contribution of FcγRs to PCV13-elicited pathogen capture by KCs. FcγRs on KCs were assessed for their contributions to cellular capture of PCV13 immune serum-opsonized bacteria by primary murine KCs (L) and bacterial capture by KCs (M) in PCV13-immunized WT, *Fcgr1^-/-^*, *Fcgr2b^-/-^*, *Fcgr3^-/-^*or *Fcgr2b/3/4^-/-^* mice. n = 3 for (L); quantitation in (M) was shown as pooled data of 6-31 FOVs from two mice in each group. Dotted line indicates the limit of detection. Data are presented as the mean ± SEM. Statistics: (C, D and K) two-way ANOVA; (E) log-rank (Mantel-Cox) test; (L and M) unpaired two-tailed Student’s t test.

The essentiality of Fc**γ**RIIB to PCV13-elicited endothelial pathogen capture was further verified by IVM imaging, which showed the EC’s ability to capture antibody-opsonized pneumococci is completely ablated in the absence of Fc**γ**RIIB *in vivo*. Specifically, PCV13-immunized *Fcgr2b^-/-^* but not *Fcgrt^-/-^* mice failed to capture pneumococci during observation (**Fig. 5F** and **5H**; **Movie S8**). Similar finding was also found with serotype-3 pneumococci (**fig. S8A-B**; **Movie S3)**. By comparison, an equivalent level of immobilized pneumococci on KCs was found between PCV13-immunized wildtype and *Fcgr2b^-/-^*mice (**Fig. 5F**; **fig. S8B**). The essentiality of Fc**γ**RIIB in EC’s recognition of IgG-opsonized was further confirmed by CPS3-specific IgG1 monoclonal antibody 5F6, which confers passive protection against serotype-3 pneumococci in a Fc**γ**RIIB-dependent manner (*40*). In contrast to its potent enhancement of EC’s bacterial capture in wildtype mice, 5F6 had no detectable effect either on the EC’s activity (**Fig. 5G** and **5I**; **Movie S6**), or on hepatic trapping of pneumococci (**fig. S8C**) in *Fcgr2b^-/-^* mice. These results showed that Fc**γ**RIIB was the sole Fc**γ**R on ECs for mediating IgG1 antibody-opsonized pneumococci, but was dispensable for PCV13-elicited bacterial capture by KCs.

Finally, we characterized the mechanism of KC-mediated capture of IgG-opsonized pathogen. Proteomic analysis detected the expression of the five known Fc receptors on murine KCs (**Fig. 5J**; **Table S2)**. Only Fc**γ**RIII significantly enhanced bacterial adhesion to CHO cells in the presence of PCV13 immune serum (**Fig. 5K)**. Furthermore, primary murine KCs from Fc**γ**RIII-deficient mice showed marked reduction in immune serum-mediated binding towards pneumococci (**Fig. 5L**; **fig. S9)**. IVM imaging also showed a significant impairment of KCs in bacterial capture in Fc**γ**RIII-deficient mice (**Fig. 5M**; **fig. S8D)**. Interestingly, primary murine KCs from Fc**γ**RIIB/III/IV-deficient mice showed even greater degree of reduction in bacteria. Collectively, these results indicate that KCs rely on multiple Fc**γ**Rs, particularly Fc**γ**RIII, to capture IgG-opsonized bacteria.

### PPV23 enables Kupffer cells to capture pneumococci by IgM-initiated complement activation

Because ECs primarily depend on IgG1 antibody and the complement system for PCV13-activated pathogen capture and killing in the liver, respectively, we assessed their potential roles in PPV23-elicited pathogen clearance. In contrast to the IgG antibodies elicited by PCV13 (**Fig. 4A**), PPV23-immunized wildtype mice produced a significant level of IgM (but not IgG) antibodies against CPS4, but no antibody was detected in vaccinated μMT mice (**Fig. 6A**). Likewise, comparing with significant bacterial clearance from the bloodstream of vaccinated WT mice at 30 min post i.v. infection, the μMT counterpart still displayed severe bacteremia similar as naïve mice (increased by 81.0 folds) (**Fig. 6B**). This result indicated that IgM antibodies are essential for PPV23-activated pathogen clearance. We next tested the role of the complement system in PPV23-induced bacteria capture and killing. Although PPV23 immunization led to a comparable level of anti-CPS4 IgM antibody in WT and *C3*^-/-^ mice (**Fig. 6A**), vaccinated *C3*^-/-^ mice exhibited 35.6-fold more bacteria in the bloodstream than the WT counterpart (**Fig. 6B**), showing that the complement system is also crucial for PPV23-elicited pathogen clearance.

**Fig. 6.**
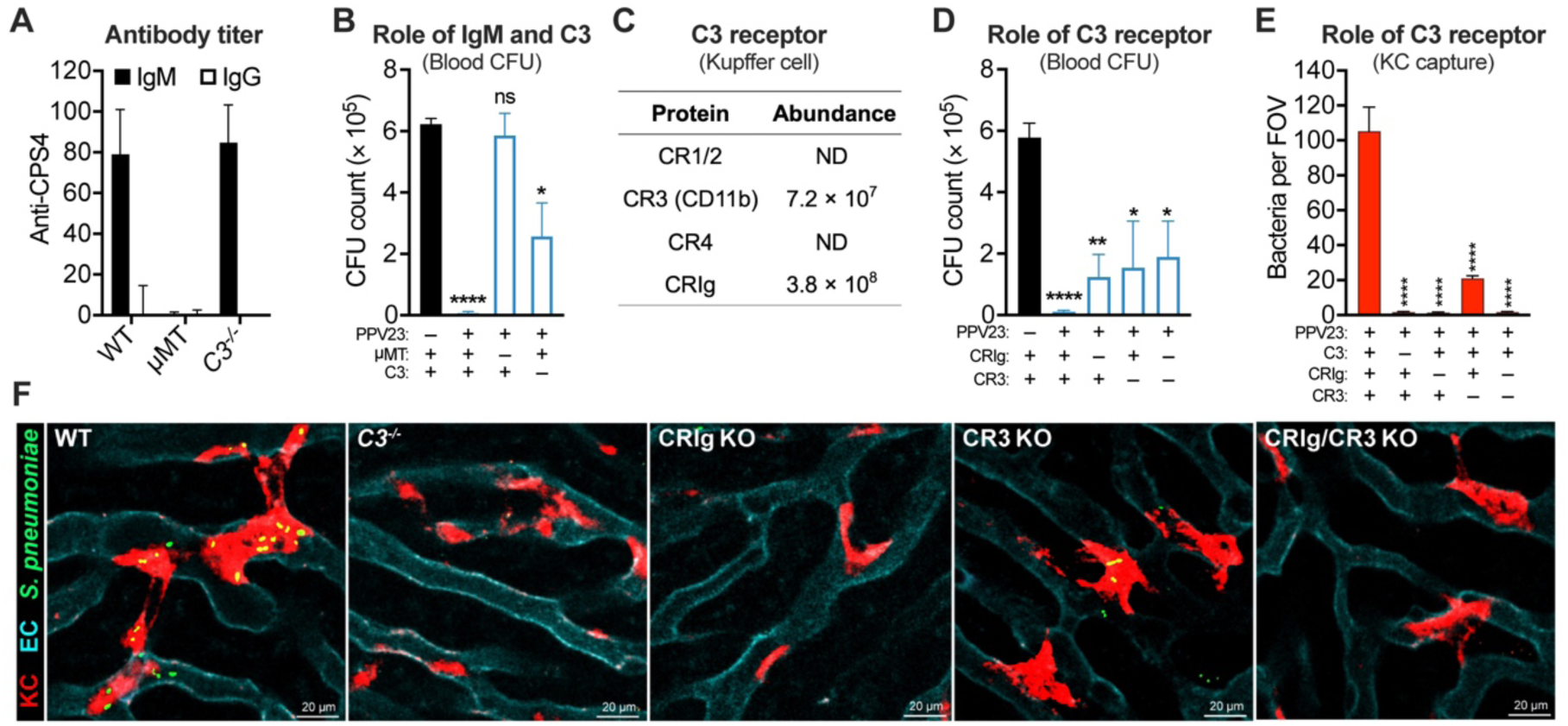
Requirement of IgM and complement system in PPV23-activated pathogen capture by Kupffer cells. (**A-E**) Dominant roles of IgM antibodies and complement system in PPV23-activated pathogen capture by KCs. Anti-CPS4 IgG and IgM antibodies were detected in sera of PPV23-immunized WT, μMT or C3-deficient (*C3^-/-^*) C57BL/6J mice by ELISA (A); n = 4-12. The roles of IgM antibodies and C3 complement in PPV23-activated pathogen clearance from bloodstream in naïve and PPV23-immunized WT (n = 4), μMT (n = 3) or *C3^-/-^* (n = 5) C57BL/6J mice are shown as the CFU count in blood (B). Expression of known C3 receptors in primary murine KCs was detected by proteomic analysis of total membrane proteins and presented as in Fig. 5A (C); The impact of complement receptors CRIg and CR3 on PPV23-activated pathogen clearance from bloodstream (D) and KCs (E) of C57BL/6J mice are assessed as in (B) and Fig. 3F, respectively; n = 3-6 in (D); quantitation in (E) was shown as pooled data of 10-20 FOVs from two mice in each group; Bacterial capture in (E) is demonstrated in (F) and **Movie S9**. The presence and absence of PPV23 immunization and individual genes are collectively expressed as + and -, respectively. (**F**) Demonstration of the essentiality of C3 and C3 receptors (CRIg and CR3) in PPV23-activated pathogen capture by KCs. Representative IVM images of the liver sinusoids are shown to demonstrate that C3 and C3 receptor (CRIg and CR3) are responsible for the KC capture of the pathogen in PPV23-immunized mice. Related data are also presented in (E) and **Movie S9**. Data shown in (A and C) were representative of duplicate experiments; data are presented as the mean ± SEM. Statistics: (B, D and E) unpaired two-tailed Student’s t test.

We further sought to determine how IgM and C3 mediate PPV23-elicited pathogen clearance by KCs since KCs are the major immune cells that execute PPV23-elicited pathogen capture in the liver (**Fig. 3L**). Three known IgM receptors (FcμR, Fcα/μR and pIgR) have been identified (*42*), but only pIgR was detected in KCs (**fig. S10A**; **Table S2**). pIgR is known for transport of IgA and IgM antibodies to mucosal surfaces, but not for pathogen clearance (*43*). In the context of the C3 importance in PPV23-elicited pathogen clearance, we hypothesized that IgM promotes pathogen capture by KCs via activating C3 on pneumococcal surface and thereby enabling C3-opsonized bacteria to be recognized by C3 receptor(s) on KCs. We thus tested the impact of C3 receptor(s) on PPV23-elicited pathogen clearance. Our proteomic analysis revealed the expression of CRIg and complement receptor 3 (CD11b subunit of CR3) by KCs, out of the five existing complement receptors (**Fig. 6C**; **Table S2**). We thus tested potential role of CRIg and CR3 in PPV23-elicited immunity using gene-deficient mice (**fig. S10B**). PPV23-immunized single KO mice of CRIg and CR3 showed a 11.8- and 14.6-fold more bacteria in the bloodstream, respectively, as compared with the WT counterpart (**Fig. 6D**; **fig. S10C-D**). Consistently, the double KO mice also showed a similar level of functional impairment. Consistent with the significance of the complement system in the PPV23-activated immunity, IVM imaging uncovered that the vaccinated *C3*^-/-^ mice completely lost the capacity of pathogen capture compared to the WT mice (**Fig. 6E-F**; **Movie S9**), which is stark contrast to the dispensable role of C3 in PCV13-induced bacterial capture (**Fig. 5H**). Furthermore, PPV23-immunized mice lacking CRIg and/or CR3 also showed significant KC impairment in bacterial capture. These results showed that C3 and its cognate receptors CRIg and CR3 on KCs were involved in PPV23-elicited pathogen capture by KCs. In summary, our data show that PPV23 achieves its vaccine efficacy through IgM-mediated activation of C3 and engagement of C3 receptors CRIg/CR3 on KCs for pathogen capture and clearance in the liver.

### The vaccine-activated immunity in the liver operates for other invasive pathogens

To determine if the Fc**γ**RIIB-mediated pathogen capture mechanisms could be applied to other vaccines, we tested conjugate vaccine MCV2 of *N. meningitidis* (meningococcus), a major pathogen of bacterial meningitis (*44*). MCV2 consists of diphtheria toxoid-conjugated CPSs of serotype A and C. MCV2-immunized mice produced significant levels of IgG antibody against both antigens (**fig. S11A**) and accelerated clearance of serotype-C meningococci from the bloodstream to the liver (**fig. S11B-D)** in the first 30 min post infection. In contrast to meningococcal capture only by KCs in the naïve mice, MCV2 immunization expanded to and enabled ECs to surpass KCs as the major hepatic cells to capture the bacteria, as manifested by dramatic increase in EC-bound bacteria in the liver sinusoids of vaccinated mice. Of note, murine KCs in the naïve mice appeared to have higher capture capacity on meningococci than pneumococci, perhaps due to preferential recognition of meningococci versus pneumococci capsule (**Fig. 3**) (*29*). Like PCV13-elicited pathogen capture by ECs (**Fig. 5H**), MCV2-elicited meningococcal capture by ECs fully relied on Fc**γ**RIIB (**Fig. 7A**; **Movie S10**). Interestingly, KCs of *Fcgr2b*^-/-^ mice maintained strong binding to serotype-C meningococci, despite lack of EC-mediated capture. This result shows that vaccine-elicited pathogen capture by ECs broadly operates beyond the pneumococcal conjugate vaccine.

**Fig. 7.**
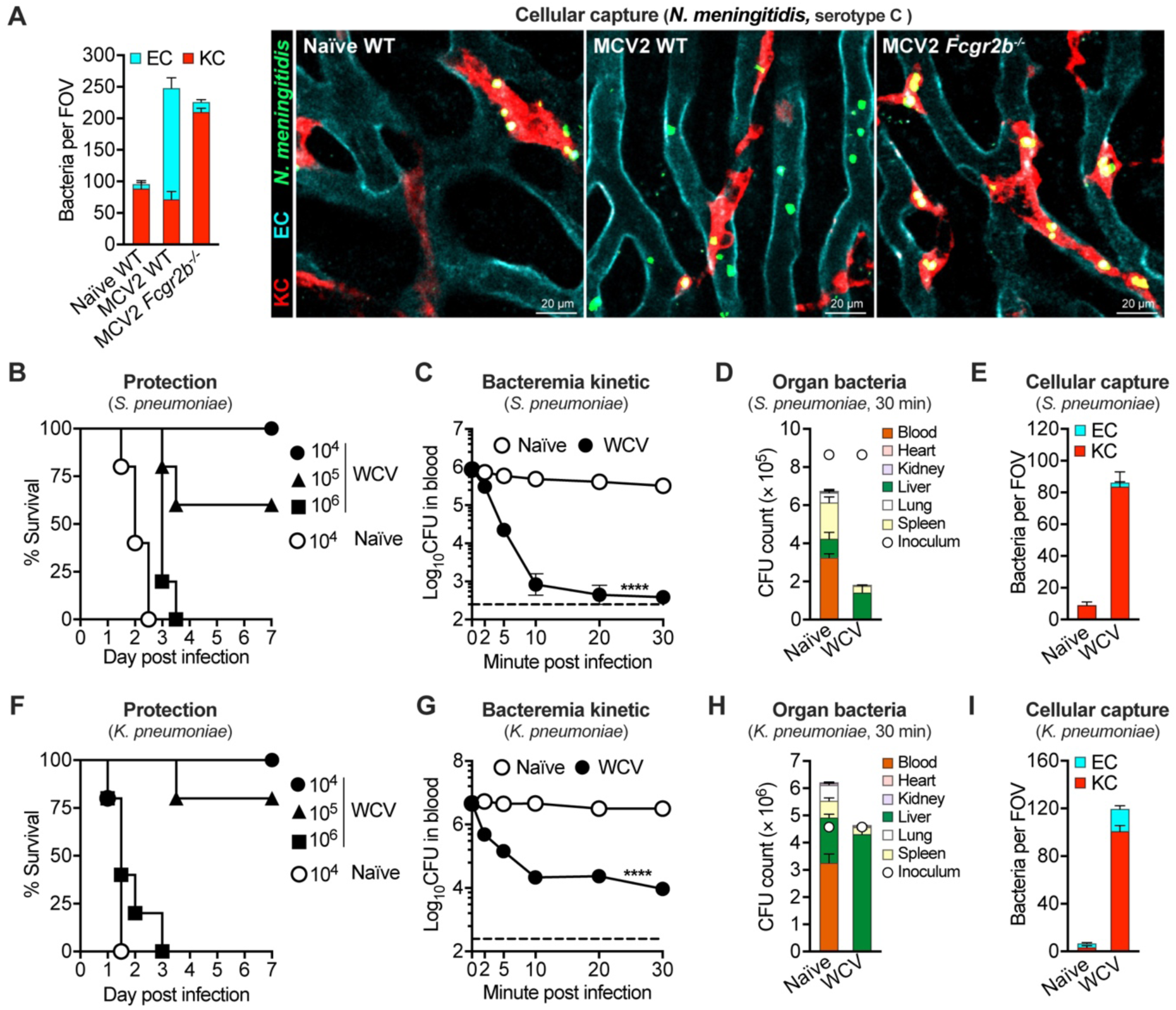
Liver-based pathogen capture activated by other vaccines. (**A**) Requirement of FcγRIIB for vaccine-activated capture of *N. meningitidis* by liver ECs. Bacterial capture was visualized and quantified by IVM imaging in naïve WT mice and MCV2-immunized WT or *Fcgr2b*^-/-^ mice post i.v. infection with 5 × 10^7^ CFU of serotype-C *N. meningitidis*; data presented as in Fig. 3L. Bacterial capture is demonstrated in **Movie S10**. Quantitation was shown as pooled data of 5-10 FOVs from two mice in each group. (**B-E**) Activation of Kupffer cell-mediated pathogen capture by whole cell vaccine of *S. pneumoniae.* Inactivated TIGR4 were used as whole cell vaccine to immunize CD1 mice and assess vaccine-elicited protection against lethal infection of TIGR4 in a 7-day period post i.p. infection (B). The short-term impact of the vaccination is reflected by early clearance (C) and hepatic capture of the pathogen post i.v. infection with 10^6^ CFU of TIGR4 (D); KC- and EC-specific capture of the pathogen (E) post i.v. infection with 5 × 10^7^ CFU of TIGR4. The experiments in B, C, D and E were carried out as in Fig. 1B, 2A, 2B and 3F, respectively. Bacterial capture in (E) is demonstrated in **Movie S11**. n = 5 and 3 (B) and (C and D), respectively; quantitation in (E) was shown as pooled data of 5 FOVs from two mice in each group. (**F-I**) Activation of Kupffer cell-mediated pathogen capture by whole cell vaccine of *Klebsiella pneumoniae.* Inactivated serotype-K1 *K. pneumoniae* strain NTUH-K2044 was used to immunize CD1 mice and assess the protection efficacy against lethal infection of K2 *K. pneumoniae* (F). The short-term impact of the vaccination is reflected by early clearance (G) and hepatic capture (H) of the pathogen post i.v. infection with 5 × 10^6^ CFU of *K. pneumoniae*; KC- and EC-specific capture of the pathogen (I) post i.v. infection with 5 × 10^7^ CFU of *K. pneumoniae*. Bacterial capture in (I) is demonstrated in **Movie S11**. n = 5 and 3 (F) and (G and H), respectively; quantitation in (I) was shown as pooled data of 9 FOVs from two mice in each group. Dotted line indicates the limit of detection; data are presented as the mean ± SEM. Statistics: (C and G) two-way ANOVA.

To determine if the liver-based pathogen capture mechanisms can be broadly applicable to other types of microbial vaccines, we further studied a non-polysaccharide vaccine against *S. pneumoniae* as illustrated in **figure S11E**. The immunization induced the high levels of IgG antibodies against *S. pneumoniae* (**fig. S11F**), and conferred a full protection against lethal challenge with 10^4^ CFU of *S. pneumoniae* (**Fig. 7B**). Vaccinated animals demonstrated effective pathogen clearance in the first 30 min after inoculation (**Fig. 7C**). Measurement of CFU in the liver revealed efficient capture during this period (**Fig. 7D**). Further characterization using IVM imaging showed that nearly all of the immobilized bacteria were bound to KCs, with few on ECs, in the liver sinusoids of immunized mice (**Fig. 7E**; **fig. S11G**; **Movie S11**). Similar results were also obtained in mice with a whole cell vaccine against *Klebsiella pneumoniae*, a major pathogen of hospital-associated blood infections (*45*), in terms of antibody production (**fig. S11H**), protection efficacy (**Fig. 7F**), bacterial clearance from the bloodstream (**Fig. 7G**), and bacteria capture by KCs in the liver (**Fig. 7H-I**; **fig. S11I**; **Movie S11**). Taken together, these results have demonstrated that vaccine-activated pathogen capture by KCs in the liver confers significant of immunoprotection against blood infections by virulent bacteria, but the KC-mediated protection efficacy is much less effective than the EC-mediated vaccine immunity.

## DISCUSSION

### The liver as the major executive organ for vaccine protection

The lack of effective vaccines has made it difficult to prevent many important human and animal diseases. One of the major obstacles to develop potent vaccines is our blurry understanding of how potent vaccines achieve more effective pathogen clearance in the process of infection. This study reveals that vaccine-activated cellular processes of pathogen capture in the liver are responsible for superior protection against blood infections of virulent bacteria by taking advantage of PCV13 and PPV23. Consistent with its superior protection in humans (*4*), PCV13 confers a superior protection to PPV23 against pneumococcal infection in mice (Fig. 1). Opposite to the paradigm that vaccine-activated immunity eliminates blood-borne bacteria by circulating phagocytes (*4, 5*), we found that virulent bacteria in blood were captured and killed in the liver sinusoids of vaccinated mice in a vaccine-specific manner (Fig. 2). In line with its superior protection, PCV13 activates a more effective program of pathogen clearance by engaging both the liver KCs and ECs, whereas PPV23 activates only KCs with a less effective rate of pathogen capture (Fig. 3). At the molecular level, PCV13 drives astonishingly rapid rate of pathogen clearance by inducing abundant IgG antibodies and thereby enhancing FcγR-mediated capture of IgG-opsonized bacteria (Fig. 4). In particular, the FcγRIIB-mediated pathogen capture by ECs is primarily responsible for the superior pathogen capture and protection of PCV13 (Fig. 5). Importantly, we have confirmed the vaccine efficacy-defining mechanisms of pathogen clearance with other invasive pathogens (Fig. 6 and 7). While the liver is shown to trap antibody- and complement-opsonized pneumococci from the bloodstream (*13, 35*), it remains unknown what cells are responsible for this hepatic activity and how it contributes to vaccine efficacy. To the best of our knowledge, this is the first time to reveal that highly effective adaptive immunity against blood-borne infections depends on vaccine-activated pathogen capture by ECs in the liver.

### The liver sinusoidal endothelial cells: the cellular basis of effective vaccines

To the best of our knowledge, this is the first observation that ECs are capable of massive capture of particles as large as bacteria (∼2 µm). KCs are well known for the ability to capture circulating bacteria and fungi (*19-22, 46, 47*). Based on the “dual-cell principle” of waste clearance (*48*), KCs take up large particles (>0.5 µm) by phagocytosis, whereas ECs preferentially eliminate soluble macromolecules and small particles (<0.3 µm) by endocytosis. ECs are also recognized as a major scavenger in the clearance of small immune complexes and antibody-opsonized viruses (*49, 50*). However, the liver ECs have not been reported to capture IgG-opsonized bacteria, which is unequivocally manifested by EC’s capture of three-quarters of total immobilized pneumococci in the liver sinusoids of PCV13-immunized mice in the early phase of blood infection. More importantly, the EC-mediated pathogen capture can be only elicited by the vaccines with superior protection (e.g. PCV13), but not by the less effective vaccines (e.g. PPV23 and whole cell vaccines). This striking feature makes the liver ECs as a cellular hallmark for highly effective vaccines.

### Advantage of IgG1 antibodies in promoting EC’s capture of pathogens

We have uncovered two critical determinants of vaccine-elicited pathogen capture by ECs in mice: 1) abundant production of IgG1 antibody, and 2) unique expression of FcγRIIB (*49*). Only KCs but not ECs were found to capture IgG-opsonized bacteria under the conditions of relatively low IgG abundance, but a shift from KCs to ECs occurred proportionally with the increase in IgG concentration. Besides the importance of the antibody abundance, the IgG1 subtype of vaccine-elicited antibodies is a prerequisite for the EC-mediated pathogen capture although the precise mechanism behind the IgG1 “favoritism” toward FcγRIIB remains to be defined. The requirement of abundant antibodies for EC-mediated pathogen capture is consistent with the abundant expression of FcγRIIB by ECs, a low-affinity FcγR (*51*). The dominant pathogen capture by ECs in PCV13-immunized mice appears to be accomplished primarily by outnumbered ECs and FcγRIIB in the liver sinusoids. The ECs constitute approximately a half of liver NPCs (*15*); the vast majority of murine FcγRIIB is expressed by liver ECs although the receptor is also found on KCs and other immune cells (*49*). Our results indicate that, although murine FcγRIIB has relatively lower affinity to IgG than other FcγRs, this “weakness” is overcome by the abundance of IgG molecules coated on bacterial cells. While previous studies have shown that IgG Fc receptors significantly affect the protective efficacy of antibodies against multiple viral pathogens (*52–54*), this work appears to be the first to demonstrate that antibody subtypes substantially affect the clearance of bacterial pathogens.

### Three mechanisms of antibody-mediated vaccine protection

This study reveals that the vaccine-elicited antibodies accomplish the clearance of blood-borne bacteria by at least three molecular mechanisms in the liver. Antibody titer has been often used as an immune correlate of vaccination against invasive bacterial diseases (*2, 55, 56*), but it remains unknown which immune cell(s) are primarily responsible for the clearance of antibody-opsonized pathogens at the sites of infection. As exemplified by PCV13, the most effective mechanism operates when pathogens are abundantly coated with IgG antibodies and subsequently captured by both the liver KCs and ECs via FcγR. While KCs are able to capture IgG-opsonized bacteria via multiple FcγRs, the FcγRIIB-mediated pathogen capture by ECs particularly defines the superior protection of PCV13. The second mechanism involves the activation of KC’s pathogen capture, but not that of ECs, by relatively low levels of IgG antibodies, which seems to be mediated by high-affinity FcγRs. This appears to be responsible for the relatively low level of immunoprotection of the *K. pneumoniae* and *S. pneumoniae* whole cell vaccines. As represented by PPV23, the third mechanism involves vaccine-induced IgM antibodies, which likely consists of three sequential steps: 1) IgM binding to bacteria, 2) immune complex-mediated activation of C3 deposited on bacteria, 3) C3-opsonized bacterial binding to CRIg and CR3 on KCs. Strict requirement for C3 in IgM-mediated pathogen capture by KCs strongly suggests that no IgM Fc receptor is involved in this mechanism.

### Two modes of the complement action in vaccine protection

Our results also indicate that the bacteria captured by ECs are effectively killed in a C3-dependent manner. Pneumococci trapped in the livers of PCV13-immunized mice were mostly eliminated in the first 30 min post i.v. infection with 10^6^ CFUs (Fig. 2). Due to EC’s dominance in hepatic pathogen capture in the vaccinated mice (Fig. 3), it is reasonable to conclude that EC-bound bacteria are effectively killed at the very early phase of septic infection. Even in the absence of major phagocytes, the vaccine-elicited hepatic trapping and killing of bacteria are still efficient, in which all the bacteria captured in the liver are bound to ECs (Fig. 2 and 3). This notion is supported by the extreme importance of FcγRIIB-mediated pathogen capture in the superior immunoprotection of PCV13 against high-dose pneumococcal infection (Fig. 5). While it remains to be determined how EC-bound bacteria are killed, our experiments in C3-deficient mice indicate that C3 is dispensable for EC’s capture of IgG-opsonized pathogen, but necessary for killing of EC-bound bacteria (fig. S6). This is in sharp contrast to the essential role of C3 in KC’s capture of IgM-opsonized bacteria (Fig. 6). Although EC-captured bacteria may be killed in an extracellular environment with the help of neutrophils or other circulating phagocytes as shown for neutrophil plucking and killing of splenic macrophage-capture pneumococci (*39*), the minimal contribution of professional phagocytes to PCV13-elicited protection efficacy has marginalized this possibility (Fig. 5E). Alternatively, EC-bound bacteria may be killed after being taken up by a C3-mediated internalization mechanism. It has been documented that, under the conditions where the phagocytic capacity of KCs is compromised, the ECs take up latex particles of 1.0 µm via typical phagocytosis (*57*). If EC-bound bacteria are taken up by C3-mediated internalization, an unknown C3 receptor(s) should be involved because we did not detect any of the five existing complement receptors by our proteomic analysis of murine ECs (Table S1).

### Practical implications of the liver-driven vaccine protection mechanisms

While the empirical approaches of vaccine development have achieved a great success in the control of certain devastating diseases (e.g. smallpox and polio), similar trials have not yielded effective vaccines for many other infections (e.g. malaria, tuberculosis, and AIDS) (*1*). Paying more attention to vaccine-elicited activities in *in vivo* pathogen clearance in vaccine design and evaluation, instead of molecular and cellular immune responses to immunization, may help break this ceiling effect in future vaccine development. In this context, the EC- and KC-mediated vaccine protection mechanisms may be used as functional markers for quantitative evaluation of the existing vaccine and future development of effective vaccines. Various aspects of pathogen shuffling from the bloodstream to the liver may be applied to the prediction of vaccine efficacy, such as pathogen clearance from the blood, pathogen enrichment in the liver, pathogen partition between ECs and KCs. Since IgG1 is the most potent subtype of IgG antibodies in driving pathogens to ECs (Fig. 4), monitoring vaccine-induced IgG1 production may be another *in vitro* parameter to assess protection-related traits, in addition to the measurement of antibody abundance and activities (e.g. opsophagocytosis and pathogen neutralization) (*58*). A highly effective vaccine is expected to elicit the production of abundant IgG1 antibodies, and thereby make invading pathogens captured predominantly by ECs in the liver at a rapid pace. These functional markers may be directly applicable to bacteria, fungi and parasites that cause systemic infections. With certain technical modifications, our findings may also be used for evaluation of vaccine candidates for viral pathogens that depend on viremia for dissemination and vector-borne transmission.

## MATERIALS AND METHODS

### Mice

All immunization and infection experiments were conducted in 5-week-old female C57BL/6J, CD1 (Vital River, Beijing, China) or sex-matched knockout (KO) mice in compliance with the guidelines of the Institutional Animal Care and Use Committee in Tsinghua University. All of the knockout mice were maintained in the C57BL/6J background. µMT mice, *C3^-/-^* mice and *Fcgr2b*^-/-^ mice were purchased from the Jackson Laboratory (Bar Harbor, Maine, USA). *Fcgr1*^-/-^, *Fcgr3*^-/-^, *Fcgr2b/3/4*^-/-^, *Fcgrt*^-/-^ mice were purchased from the Model Organisms Center (Shanghai, China). CRIg KO (*Vsig4*^-/-^) mice (*19*) were acquired from Genentech (CA, USA). All mice were housed as groups of 4 to 6 individuals per cage and maintained on a 12-h light-dark cycle at 22-25°C under specific-pathogen free conditions.

CRIg/CR3 double KO mice were obtained by cross breeding of CRIg KO mice (*Vsig4*^-/-^) and CR3 KO (*Itgam*^-/-^) mice. CR3 KO (*Itgam*^-/-^) mice were generated by CRISPR/Cas9 (*59*). Briefly, single guide RNA (sgRNA) (GCAAAGGCTGTTAACCAGAC) was designed to target the exon 3 of *Itgam*, which encodes the CD11b subunit of CR3, using CRISPOR (*60*). sgRNA and Cas9 were transcribed into RNA *in vitro* using MEGAshortscript Kit (Ambion) and mMESSAGE mMACHINE T7 Ultra Kit (Ambion) according to the manufacturer’s instructions. A 20-μl mixture solution containing Cas9 mRNA (100 ng/μl) and sgRNA (50 ng/μl) was microinjected into zygotes of C57BL/6J mice. Then, the injected zygotes were transferred to C57BL/6J females as described previously (*59*). The genotype of F1 offspring was based on the sequence of exon 3. The offspring with frameshift mutation (one base insertion between C42 and A43 of exon 3) of CD11b was chosen to back cross with wildtype C57BL/6J mice more than 7 generations. The knockout effect of CD11b mice was confirmed by PCR. The mice strains used in this study were listed in **Table S3**.

### Primary cells and cell lines

Primary murine liver sinusoids endothelial cells (ECs) and Kupffer cells (KCs) were isolated from fresh liver sections by magnetic-activated cell sorting (MACS) or fluorescence activated cell sorting (FACS), respectively, as described (*61, 62*). HEK293T cells were grown in DMEM medium (Gibco) with 10% fetal bovine serum (FBS) (Biological Industries), 1 mM sodium pyruvate (Corning), 1 × non-essential amino acid (NEAA) (Gibco) and 1 × penicillin/streptomycin (Corning). HEK293 suspension culture cells (Expi293F) were grown in SMM 293-TII medium (Sino Biological) with shaking of 120 rpm. CHO cells were cultured in F-12K medium (Gibco) with 10% FBS, 1 mM sodium pyruvate, 1 × NEAA, 1 × penicillin/streptomycin, 5 μg/ml puromycin (Invivogen), and/or 300 μg/ml zeocin (Invivogen). All cells were cultured at 37°C with 5% CO_2_.

### Bacterial cultivation

Pneumococcal strains were cultured in Todd-Hewitt broth (Oxoid) with 0.5% yeast extract (THY) or on tryptic soy agar (TSA, BD Difco) plate supplemented with 5% defibrinated sheep blood (FeiMoBio, China) as described (*63*). 5 μg/ml for gentamycin (Sigma-Aldrich) was added when necessary. *S. aureus* strains were cultured in tryptic soy broth (TSB, BD Difco) (at 37°C, with shaking) as described (*64*). The meningococcal strain was cultured in brain heart infusion (BHI, BD Difco) broth (at 37°C, with shaking) or BHI agar (10%) plate as described (*65*). All bacterial strains used in this study are described in **Table S4**.

### Vaccination and immuno-protection

All the vaccines used in this study were approved by the Chinese National Medical Products Administration (NMPA). Immunization with PCV13 (Minhai Biotechnology, China, NMPA S20210036), PPV23 (Minhai Biotechnology, China, NMPA S20180009) and MCV2 (CanSinoBio, China, NMPA S20210018) was carried out essentially as described (*38*). Briefly, vaccines were inoculated by intramuscular injection (50 µl/mouse, 1/10 of dose in human), and boosted twice by the same procedure with an interval of 14 days. PCV13 vaccine dilution was carried out by various dilutions with PBS in a total injection volume of 50 µl. For immunization of whole cell vaccine, 10^8^ CFU of *S. pneumoniae* TIGR4 or *K. pneumoniae* NTUH-K2044 were suspended in PBS with 0.5% formalin and incubated at 4°C overnight. Mice were immunized by subcutaneous injection of formalin-fixed bacteria mixed with 25 µl of Adju-Phos^®^ adjuvant (InvivoGen) in a total volume of 500 µl, and boosted once by the same procedure with an interval of 7 days (*66*). Immunized mice were used for immune serum collection by retro-orbital bleeding 14 days or for septic infection 3 weeks after final immunization.

Immunoprotection was assessed by intravenous (i.v.) or intraperitoneal (i.p.) injection of bacteria in mice as described (*38*). The animals were periodically sampled for blood bacteria by every 12 hrs by retro-orbital bleeding and observed for survival in a 7-day period. LD_50_, lethal dose 50%, was calculated by infection dose and percent of survival using LD_50_ calculator (Quest Graph^TM^, AAT Bioquest, Inc.). Passive protection of immune serum against lethal infection of *S. pneumoniae* was evaluated as described (*67*). Briefly, sera from naive (natural) or vaccinated (immune) mice were incubated at 56°C for 30 min for heat inactivation of complement protein (*68*). Then heat-inactivated serum (10 μl) was pre-incubated with a lethal dose of *S. pneumoniae* (10^6^ CFU in 90 μl) for 5 min before being i.v. inoculated into mice.

### *In vivo* bacterial clearance

Bacterial clearance from the bloodstream was assessed by i.v. inoculation with *S. pneumoniae* (10^6^ or 10^8^ CFUs in 100 µl PBS), *N. meningitidis* (2 × 10^8^ CFUs in 100 µl PBS) or *K. pneumoniae* (10^6^ CFUs in 100 µl PBS), and retroorbital bleeding/CFU plating as described (*69*). Bacterial 50% clearance time (CT_50_) was calculated by nonlinear regression analysis of bacteremia kinetic for individual strains using the formula CT_50_ = ln[(1-50/Plateau)/(-K)], in which Plateau and K were generated by One-phase association of the clearance ratio using GraphPad Prism version 8.3.1. CT_50_ was designated as 30 min if less than half of the inoculum was cleared from the blood at 30 min.

Viable bacteria in the liver, spleen, lung, kidney, and heart were determined by agar titration of the tissue homogenates and represented as CFUs per organ. Total bacterial burden in each animal was calculated as the summary of CFU values in the blood and major organs. The impact of immune serum and monoclonal antibodies on *in vivo* bacterial clearance was carried out similarly. The heat-inactivated serum or monoclonal antibodies was pre-incubated with a lethal dose of *S. pneumoniae* (10^6^ CFU in a total volume of 100 μl) for 5 min before being i.v. inoculated into mice.

### Immune cell depletion

Macrophages were depleted by i.v. administration of 1 mg of clodronate liposomes (CLL, Liposoma) 24 h prior to infection as described previously (*70*). Neutrophils and inflammatory monocytes were depleted by i.p. injection of 500 μg anti-Ly6G/Ly6C antibody (Gr1, BioXCell) 24 h prior to infection as described previously (*71*). The same dose of isotype-matched IgG2b (BioXCell) was used as a negative control. For detection of survival rate after depletion, 1mg of CLL was delivered through i.v. injection on day −1, 1, 3 and 5 of infection and 500 μg of anti-Ly6G/Ly6C antibody (Gr1) was administered through i.p. injection on a daily basis starting 1 day before infection until the end of the experiments. After the immune cell depletion, mice were i.v. infected with *S. pneumoniae* (10^6^ CFUs in 100 µl PBS) or *S. aureus* (10^8^ CFUs in 100 µl PBS) and observed for blood bacteria every 12 hrs and survival in a 7-day period.

The depletion efficiencies were verified by flow cytometry. Briefly, mouse whole blood cells, spleen cells and liver non-parenchymal cells (NPCs) were stained for 20 min with APC-Cy7 anti-CD45 (1:200), APC anti-CD31 (1:200), BV605 anti-CD11b (1:500), PE anti-F4/80 (1:200), FITC anti-Ly6G (1:500) and eF450 anti-Ly6C (1:500). Single cell populations were gated as neutrophils (CD31^-^CD45^+^CD11b^+^Ly6G^high^/Ly6C^+^), inflammatory monocytes (CD31^-^CD45^+^CD11b^+^Ly6G^-^ Ly6C^high^), KCs (CD31^-^CD45^+^CD11b^low^F4/80^+^), spleen red pulp macrophages (CD31^-^ CD45^+^CD11b^low^F4/80^+^). The antibodies used in this study were listed in **Table S3**.

### Enzyme-linked immunosorbent assay (ELISA)

Antibody titers after vaccination were quantified by enzyme-linked immunosorbent assay (ELISA) as described (*38*). Briefly, 96-well plates were coated with 100 µl of PBS containing 5 µg/ml capsular polysaccharide overnight at 4°C. After being blocked with 5% non-fat dry milk (BD Difco) in PBS, serially diluted mouse serum in PBS was added and incubated at 37°C for 2 h. Bound antibody was detected with goat anti-mouse horseradish peroxidase (HRP)-conjugated IgG (1:2,000) at 37°C for 1 h to detect for IgG detection or goat anti-mouse horseradish peroxidase (HRP)-conjugated IgM (1:2,000) for IgM detection. For the detection of immunoglobulin subtypes, phosphatase-conjugated IgG1-, IgG2b-, IgG2c- and IgG3-specific antibodies (Proteintech) were used. After addition of the TMB substrate (TIANGEN, China) for 5 minute-incubation, the reaction was stopped with 1 M phosphoric acid solution and the optical absorbance at wavelength of 450 nm was determined on a BioTek Synergy H1 microplate reader (BioTek).

### Recombinant murine monoclonal antibody generation

The recombinant anti-CPS3 monoclonal antibodies of different murine IgG subtypes were generated as described with minor modification (*72*). The coding sequences of variable region for light chain (VL, GenBank: DQ228148.1) and heavy chain (VH, GenBank: DQ228147.1) of anti-CPS3 monoclonal antibody 5F6 were synthesized according to the sequences previously published(*40*). The coding sequences of constant region for light chain (CL) and heavy chain (CH) of different murine IgG subtypes were synthesized as following: CL (GenBank: MH208239.1), CH_IgG1_ (GenBank: MH208238.1), CH_IgG2b_ (GenBank: LC052323.1), CH_IgG2c_ (GenBank: LC037230.1) and CH_IgG3_ (GenBank: MT019336.1). The DNA fragments of heavy chain and light chain were produced by fusion PCR with VH and CH, or VL and CL, respectively, and cloned into expression vectors (H vector for expression of heavy chain and L vector for expression of light chain) to produce full murine IgG antibodies with different subtypes. The relevant primers and plasmids were listed in **Table S4**. Antibody production was conducted by co-transfection of the heavy and light chain expression vectors into Human HEK293 suspension culture cells (Expi293F) using polyethylene mine (Sigma). Cells were subsequently cultured for 4 days before being pelleted by centrifugation. The supernatant was harvested and filtered through the 0.2 μm filter for monoclonal antibody purification.

### IgG purification

The total IgG was purified from murine serum using Protein G Sepharose 4 Fast Flow columns (GE Healthcare) according to the manufacturer’s instructions. The recombinant monoclonal antibodies of IgG subtypes were purified after incubation of filtered Expi293F cell supernatant with Protein G resin beads (GenScript) at 4°C overnight. Then the purification was processed according to the manufacturer’s instructions. Fractions containing IgG were identified by SDS-PAGE under reducing/non-reducing conditions and concentrated using Centrifugal Filters (Millipore). Protein concentration was quantified with the BCA Assay Kit (Beyotime, Shanghai, China). The IgG activity in promoting *in vivo* bacterial clearance was assessed by incubating 10^6^ CFU pneumococci with purified IgG or IgG1 isotype control (BioXcell) in 100 µl PBS at room temperature for 5 min before i.v. injection of the mixture in naïve mice.

### Intravital microscopy (IVM)

IVM imaging of mouse liver was carried out with inverted confocal laser scanning microscope as described (*70*). The liver sinusoid endothelial cells (ECs), Kupffer cells (KCs) and neutrophils were labelled by i.v. injection of 2.5 μg of AF594 anti-CD31 (ECs), AF647 anti-F4/80 (KCs) and PE anti-Ly6G (neutrophils) antibodies, respectively, 30 min before i.v. inoculation of FITC-labeled bacteria. FITC labeling was accomplished by incubating bacteria (5 × 10^7^ CFU in 100 μl PBS) with 20 µg FITC (Sigma-Aldrich) at room temperature for 30 min as described (*73*). For serum pre-treated pneumococci, 2 × 10^7^ CFU of bacteria were premixed with natural or immune serum for 30 min at room temperature before i.v. injected into naïve mouse. For monoclonal antibody-mediated pathogen capture, mice were i.v. inoculated with antibody in 100 μl of PBS immediately after i.v. injection of 5 × 10^7^ CFU of serotype-3 pneumococci.

Intravital imaging of the liver microvasculature was acquired on an inverted Leica TCS-SP8 confocal microscope using a 10×/0.45 NA and 20×/0.80 NA HC PL APO objectives. Photomultiplier tubes (PMTs) and Hybrid Photodetectors (HyD) were used to detect different pixel sizes signals (600 × 600-pixel sizes for time-lapse series and 1024 × 1024-pixel size for figures). Four laser excitation wavelengths (488, 535, 594 and 647 nm) were used by white light laser (1.5 mw, Laser kit WLL2, 470-670 nm). Real-time imaging was monitored immediately for at least 1 min after infection by LAS X Core module (version 3.7.3). At least five random fields of views (FOV) during 10-15 min post infection were chosen to enumerate bacteria per FOV. The percentage of cell-associated bacteria was calculated by dividing the bacteria captured by each cell with the total bacteria captured in the same FOV.

### Isolation of liver non-parenchymal cells

Liver non-parenchymal cells (NPC) were isolated by a combined ‘collagenase–DNase’ perfusion method as described with minor modification (*74*). Briefly, mouse was euthanized with 300 mg/kg avertin and the liver was perfused *in situ* from the portal vein with 5 ml of digestion buffer (HBSS (Corning) with 0.5 mg/ml collagenase IV (Sigma-Aldrich), 20 μg/ml DNase I (Roche) and 0.5 mM CaCl_2_ (Amresco)). The liver was then excised, gently minced to small pieces, and further digested with 10 ml of digestion buffer at 37°C for 30 min at 300 rpm. After digestion, the liver homogenate was filtered through a 70-µm cell strainer (Biologix), pelleted by centrifugation at 300 g for 5 min at 4°C, and washed twice with 10 ml ice-cold HBSS. Residual erythrocytes were lysed with 1 ml RBC lysis buffer (BioLegend) on ice for 1 min before the reaction was terminated with 10 ml ice-cold HBSS. Cell debris and hepatocytes were removed by centrifugation at 50 g for 2 min at 4°C. Liver NPCs from top aqueous phase were collected by centrifugation at 300 g for 5 min at 4°C. The enriched NPCs allow for cell type-specific analysis and/or further separation of ECs and KCs.

### Separation of ECs and KCs

Murine ECs and KCs were isolated from liver NPCs as described (*61, 62, 74*). ECs were purified by magnetic-activated cell sorting (MACS) using anti-CD146 MicroBeads (Miltenyi Biotec) according to the manufacturer’s instructions. KCs were obtained by fluorescence activated cell sorting (FACS). Briefly, every 1×10^6^ liver NPCs were stained for 20 min with APC-Cy7 anti-CD45 (1:200), APC anti-CD31 (1:200), BV605 anti-CD11b (1:500) and PE anti-F4/80 (1:200) (All BioLegend). KCs were gated as the CD45^+^CD31^-^CD11b^low^F4/80^+^ population using FACSAria SORP (BD Biosciences). Cell type-specific analysis of liver NPCs and isolated ECs and KCs were determined by flow cytometry.

### Protein mass spectrometry

The proteome of isolated murine KCs or ECs was assessed by quantitative protein mass spectrometry using membrane protein-enriched KCs or EC lysates as described (*75*). The membrane proteins of KCs or ECs were enriched using the Mem-PER Plus Membrane Protein Extraction Kit (Thermo Scientific) with Halt Protease and Phosphatase Inhibitor Cocktail (Thermo Scientific) according to the manufacturer’s instructions. Protein concentration was quantified with the BCA Assay Kit. Each 40 μg of protein samples were loaded on SDS-PAGE gel to excise protein bands for in-gel digestion and subsequent LC-MS/MS analysis using Thermo-Dionex Ultimate 3000 HPLC system combined with the Thermo Orbitrap Fusion mass spectrometer. The spectra from each LC-MS/MS run were searched against the mouse Unreviewed TrEMBL FASTA database (release 2020_05) using Proteome Discovery searching algorithm (version 1.4).

### Expression of murine proteins in CHO cells

Expression of murine proteins in CHO cells was achieved using pCDH and/or pFUGW vectors as described (*76*). The cDNA pool was generated with total RNA extracted from isolated KCs or ECs using Maxima H Minus First Strand cDNA Synthesis Kit (Thermo Scientific). The cDNAs of *Fcgr1*, *Fcgr3*, *Fcgr4* and *Fcgγ* (the common *γ* chain) were amplified with primer pairs Pr18178/Pr18179, Pr16960/Pr16961, Pr18180/Pr18181 and Pr16966/Pr16967 from the pool of KCs, respectively. The cDNAs of *Fcgr2b*, *Fcgrt* and *belat2mG* (*B2m*) were amplified with primer pairs Pr16958/Pr16959, Pr16962/Pr16963 and Pr16964/Pr16965 from the pool of ECs, respectively. Then the amplicons were cloned in the XbaI/EcoRI site of pCDH (*Fcgr1*, *Fcgr2b*, *Fcgr3*, *Fcgr4* and *Fcgrt*) or the XbaI/NheI site of pFUGW (*Fcgγ* and *B2m*). A sequence encoding the His tag at the carboxyl terminus of resulting protein was added in each construct. The relevant primers and plasmids were listed in **Table S4**.

Each of the recombinant plasmids was transfected into HEK293T cells together with lentiviral packaging plasmids pMD2.G and psPAX2 (gifts from Didier Trono, Addgene) using Lipofectamine 2000 (Invitrogen). The cell supernatants containing lentiviral particles were harvested after 48 h and filtered using 0.45 μm filter (Millipore). pCDH-lentivirus were used to transfect CHO cells with 8 μg/ml polybrene to express FcγRI (pCDH::*Fcgr1*), FcγRIIB (pCDH::*Fcgr2b*), FcγRIII (pCDH::*Fcgr3*), FcγRIV (pCDH::*Fcgr4*) and FCGRT (pCDH::*Fcgrt*). The cells were selected at the presence of 5 μg/ml puromycin for 7 days. For co-expression, the secondary pFUGW::*Fcgγ*-lentivirus for co-expression of γ chain was further transduced into CHO cell lines expressing FcγRI, FcγRIII and FcγRIV, respectively, while pFUGW::*B2m*-lentivirus for co-expression of b2m subunit was further transduced into CHO cell lines expressing FCGRT. Then, these co-expressed cell lines were selected under 300 μg/ml zeocin (Invivogen) for 10 days. The resulting cells were tested for expression of target proteins by Western blotting using mouse anti-His tag (1:5,000) and goat anti-mouse horseradish peroxidase (HRP)-conjugated secondary antibody (1:10,000), with β-actin as the loading control detected by anti-β-actin antibody (1:1,000) as described (*77*).

### *In vitro* bacterial adhesion

Bacterial adhesion to primary murine cells and CHO cells expressing IgG receptors was evaluated essentially as described (*78*). Briefly, primary ECs and KCs were resuspended in RPMI 1640 medium (Corning) with 5% FBS and 1 × penicillin/streptomycin (*79*). ECs were plated into 40 µg/cm^2^ collagen I (Corning)-coated 96-well cell culture plates while KCs were plated into 96-well cell culture plates without collagen coating as described with minor modification (*61*). CHO transfectants were prepared in the 48-well cell culture plates. At the time of testing, complete media was replaced with basic RPMI 1640 (Corning) (for ECs and KCs) or F-12K (Gibco) (for CHO) without serum or antibiotics. Bacteria at a multiplicity of infection (MOI) of 1:1 were added to each well in the presence of 10% (v/v) heat-inactivated immune or natural serum, centrifuged at 500 g for 5 min and incubated at 37°C with 5% CO_2_ for 30 min. Free bacteria were determined by agar titration of the supernatants; cell-associated bacteria were enumerated by CFU plating after the cells were lysed using ice cold sterile H_2_O. Finally, the bacterial binding percent was calculated by dividing cell-associated CFUs with the total CFUs.

### Quantification and statistical analysis

All experiments were performed with at least two biological replicates except animal experiments with minimal sample size that determined based on the animal welfare guidelines. All data are presented as Mean ± SEM. For comparisons between two groups, analysis was done by two-sided unpaired *t* test. For more than two groups, two-way ANVOA multiple comparisons test was used. Flow cytometry data were analyzed using FlowJo (version 10.4). Statistical analyses were performed using GraphPad Prism software (version 8.3.1). A *P* value of <0.05 was considered significant (ns, nonsignificant; \**P* < 0.05, \*\**P* < 0.01, \*\*\**P* < 0.001, and \*\*\*\**P* < 0.0001).

## Acknowledgments

We thank Wanli Liu for the CHO cells, and the Tsinghua research platforms for assistance in animal experimentation (Laboratory Animal Research Center), flow cytometry and IVM imaging (Center for Cell Biology), and protein mass spectrometry (Center for Proteomics).

## Author contributions

Conceptualization: J.W., H.A., L. Z., J.-R.Z.;

Experimentation: J.W., M.D., H.A., Y.L., S.W., Q.J. H.D.;

Methodology: J.W., M.D., H.A., Y.L., S.W., Q.J., H.D.;

Investigation: J.W., M.D., H.A., J.Z., D.E.B., H.Z., L. Z., J.-R.Z;

Material: J.Li., T.Z., J.Z., J.Liu, H.Z., Z.S.;

Funding: J.W., J.-R.Z.;

Project administration: J.-R.Z.;

Supervision: J.W., J.-R.Z.;

Writing original draft: J.W., H.A., M.D., H.Z., L. Z., J.-R.Z.

## Data and materials availability

All data are available in the main text or the supplementary materials.

## Supplementary Materials

### SUPPLEMENTAL FIGURES

**Fig. S1.**
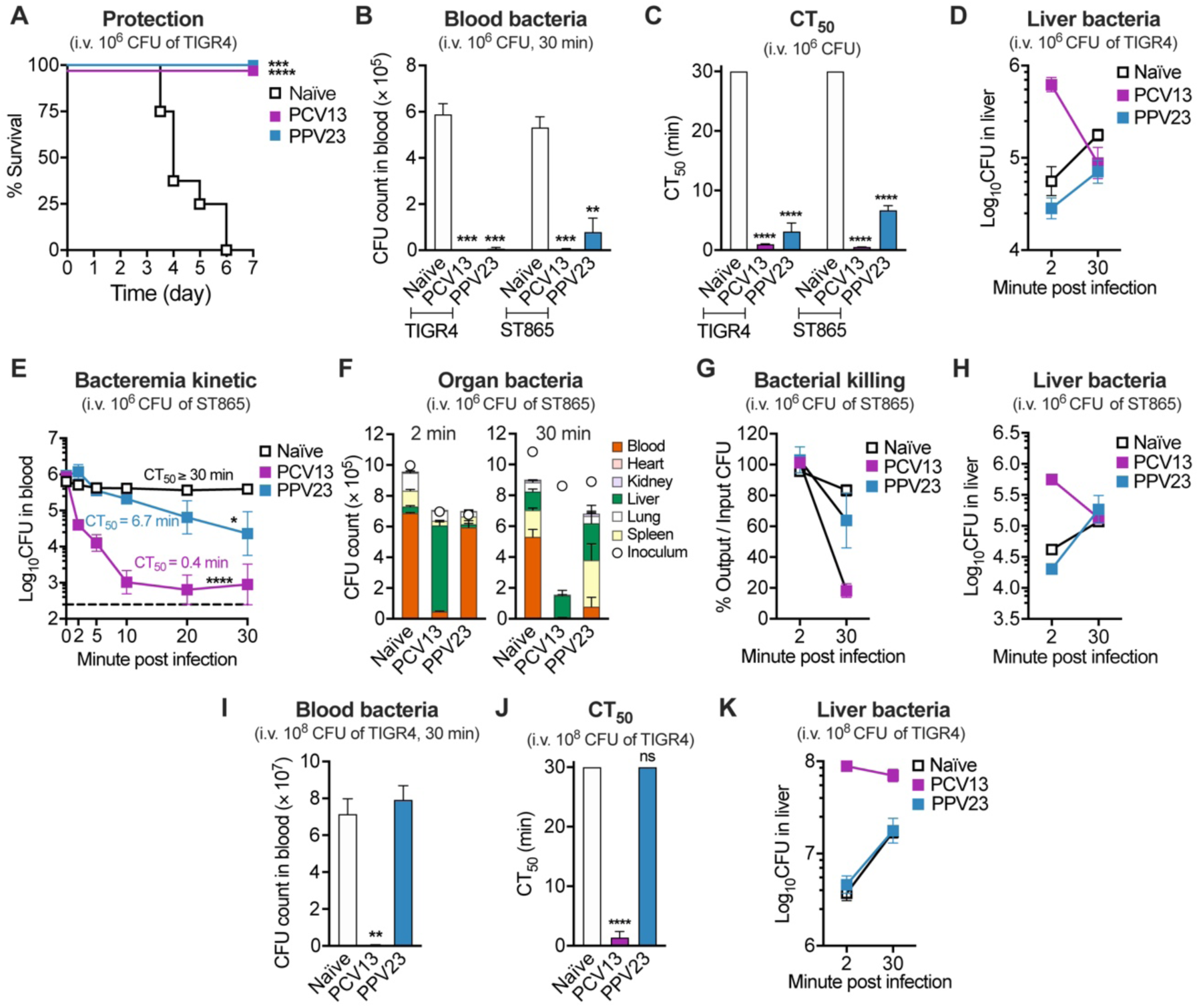
Faster capture and killing of pneumococci in liver of PCV13- than PPV23-immunized mice. (**A**) Survival of PCV13- or PPV23-immunized mice were compared with naïve mice after i.v. infection with 10^6^ CFU of TIGR4. n = 5-8. (**B** and **C**) Bacterial CFU in blood (B) and CT_50_ (C) of naïve or PCV13- and PPV23-immunized mice after i.v. infection with 10^6^ CFU of TIGR4 or serotype-3 ST865 pneumococci. CT_50_ represents the time for clearing 50% of the inoculum from the circulation. n = 3. (**D**) Viable bacteria in liver of each mouse at 2 and 30 min, which reflects *in vivo* bacterial killing process in liver during the first 30 min post infection with 10^6^ CFU of TIGR4. n = 3. (**E**-**H**) Vaccine-activated pathogen clearance by liver of low-dose serotype-3 pneumococci. Mice were immunized and subsequently infected with 10^6^ CFU of serotype-3 pneumococci ST865 to determine bacteremia kinetic (E), organ bacteria (F), total bacterial killing (G) and bacterial killing in liver (H). n = 3. (**I-K**) Bacterial CFU in blood (I), CT_50_ (J) and bacterial killing in liver (K) of naïve or PCV13- and PPV23-immunized mice after i.v. infection with 10^8^ CFU of TIGR4. n = 3. Data are presented as the mean ± SEM; dotted line indicates the limit of detection. Statistics: (A) log-rank (Mantel-Cox) test; (B, C, I and J) unpaired two-tailed Student’s t test; (E) two-way ANOVA.

**Fig. S2.**
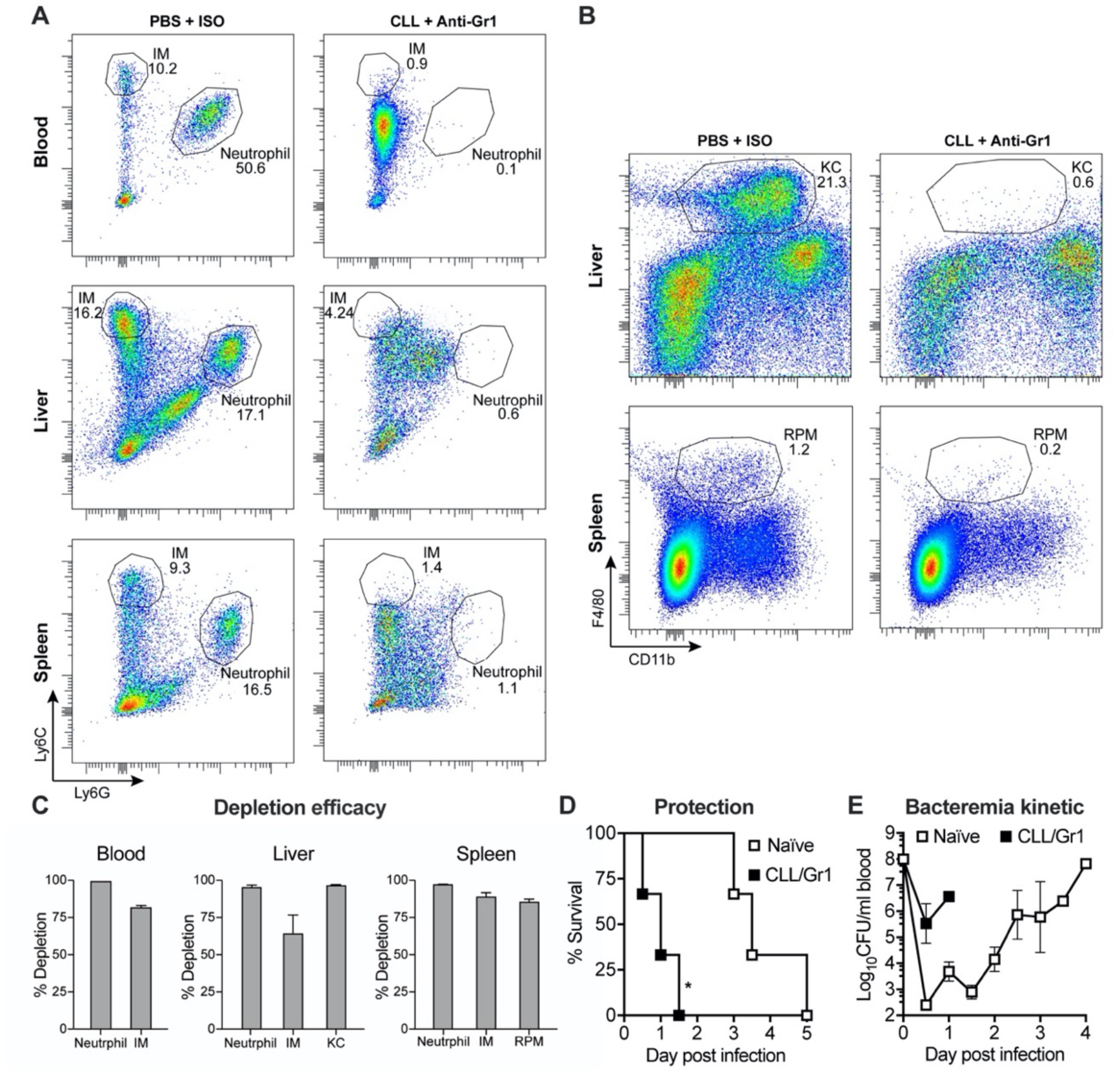
Depletion of major phagocytes in blood, spleen and liver. (**A-C**) Depletion efficiency of neutrophils, monocytes and macrophages verified by flow cytometry. Mice were treated with clodronate liposomes (CLL) and anti-Ly6C/Ly6G (Gr1) or PBS and isotype control (ISO) 24 hr before detection. The ratios of neutrophils (Ly6G^high^/Ly6C^+^) and inflammatory monocytes (Ly6G^-^/Ly6C^high^) were quantified in the myeloid cells (CD31^-^/CD45^+^/CD11b^+^) (A); the ratios of liver Kupffer cells (KCs) (CD11b^low^/F4/80^+^) and spleen red pulp macrophages (RPMs) (CD11b^low^/F4/80^+^) were measured in the immune cells (CD31^-^/CD45^+^) (B). Quantitation of depletion efficiency of major phagocytes in the blood, liver and spleen (C). n = 3. (**D** and **E**) Importance of major phagocytes (macrophages, neutrophils and monocytes) in pathogen clearance of *Staphylococcus aureus*. The survival (D) and bacteremia kinetic (E) of naïve or CLL/Gr1-treated mice were recorded after i.v. infection with 10^8^ CFU of *S. aureus*. n = 3. Data shown in (C) were representative of duplicate experiments; data are presented as the mean ± SEM. Statistics: (D) log-rank (Mantel-Cox) test.

**Fig. S3.**
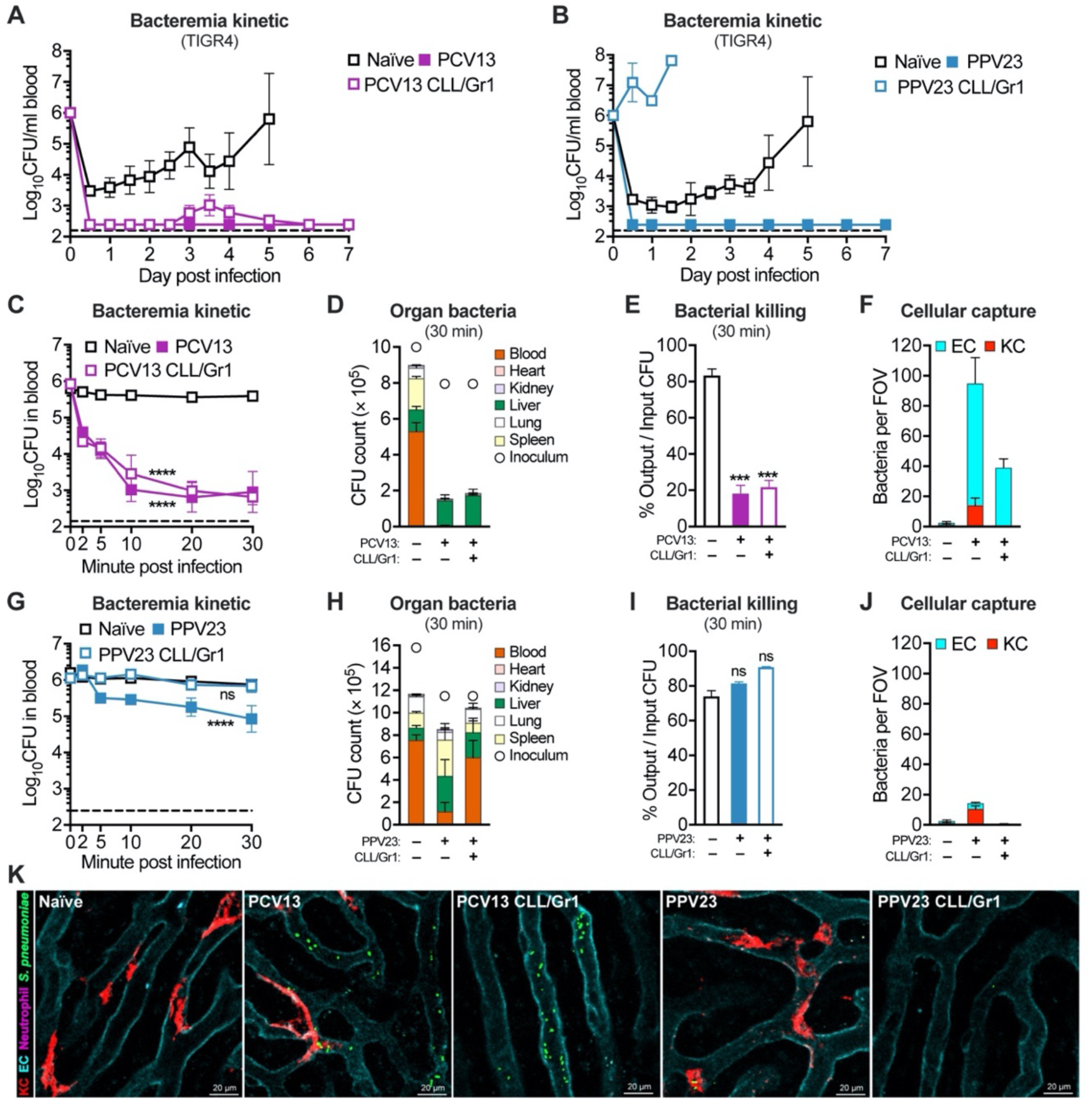
PCV13 and PPV23 activate distinct cellular mechanisms of pathogen capture in the liver. (**A** and **B**) Bacteremia kinetic of naïve and PCV13- (n = 7-8) (A) or PPV23-immunized (n = 5) (B) normal or phagocytes-depleted (CLL/Gr1) mice during the first 7 days after i.v. infection with 10^6^ CFU of TIGR4. (**C-F**) Dispensability of major phagocytes in PCV13-elicited immunity. Naïve and immunized mice without (PCV13) or with (PCV13 CLL/Gr1) CLL/Gr1 treatment were i.v. infected with ST865 as in Fig. 3A. Bacteremia kinetic (C), organ bacteria (D), bacterial killing (E) and bacterial capture by ECs and KCs (F) are presented as in Fig. 3C, 3D, 3E and 3F, receptively. The process of bacterial capture is demonstrated in (K) and **Movie S3**. n = 3 for (C-E); quantitation in (F) was shown as pooled data of 5 FOVs from two mice in each group. (**G-J**) Essentiality of major phagocytes in PPV23-elicited immunity. Bacteremia kinetic (G), organ bacteria (H), bacterial killing (I) and bacterial capture by ECs and KCs (J) in PPV23-immunized normal (PPV23) and phagocytes-depleted (PPV23 CLL/Gr1) mice post i.v. infected with ST865 as in Fig. 3A are presented as in (C-F). The process of bacterial capture is demonstrated in (K) and **Movie S3**. n = 3 for (G-I); quantitation in (J) was shown as pooled data of 5-20 FOVs from two mice in each group. (**K**) Representative *in vivo* real-time imaging of vaccine-activated pathogen capture in the liver sinusoids. Serotype-3 pneumococci ST865 (green), Kupffer cells (KCs, red), sinusoidal endothelial cells (ECs, cyan) and neutrophils (magenta) were visualized in naïve and immunized mice without or with depletion of major phagocytes by CLL/Gr1 treatment as illustrated in Fig. 3A. Data are presented as the mean ± SEM; dotted line indicates the limit of detection. Statistics: (C and G) two-way ANOVA; (E and I) unpaired two-tailed Student’s t test.

**Fig. S4.**
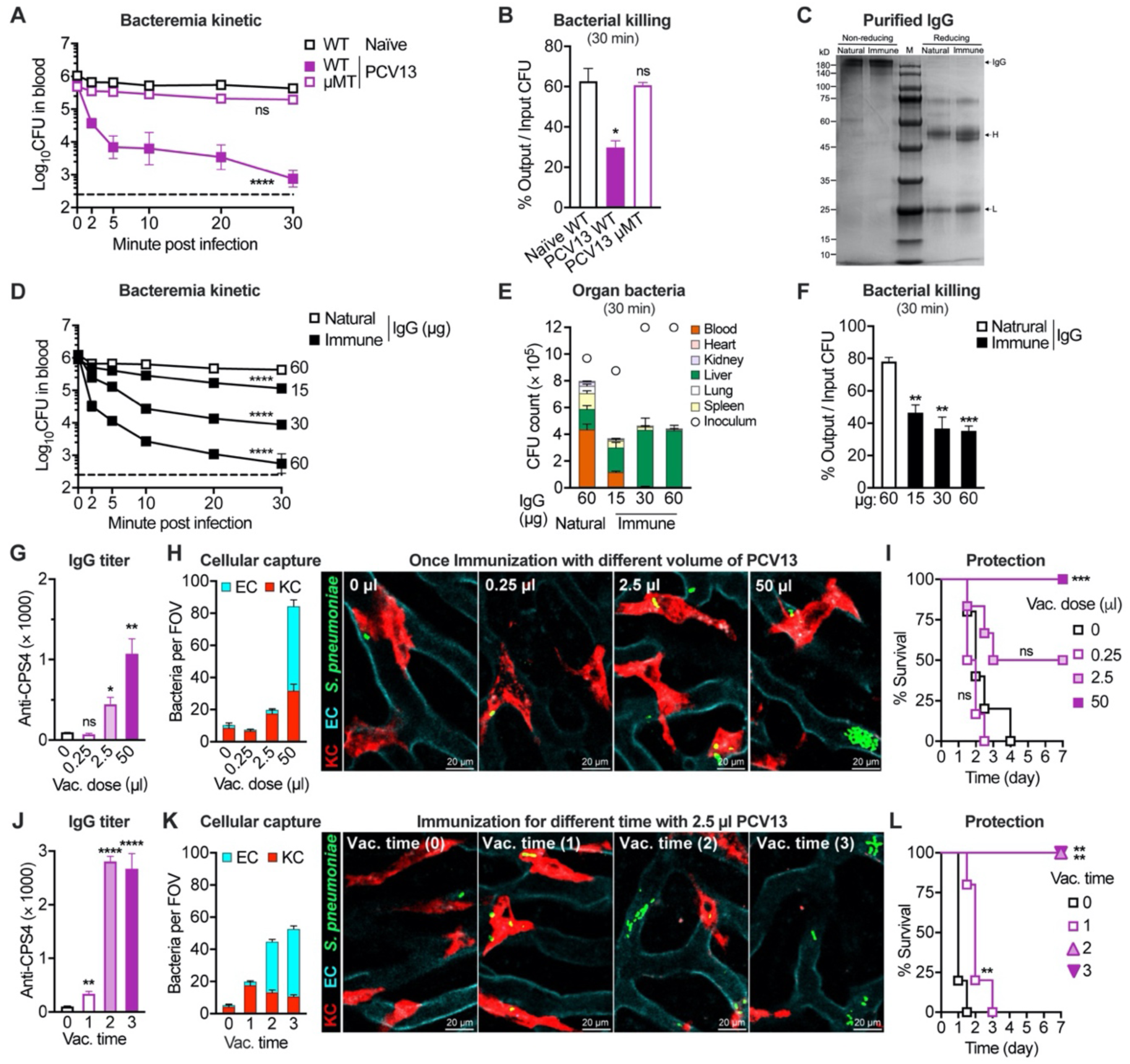
The antibody abundance-dependent endothelial capture of pneumococci. (**A** and **B**) Bacteremia kinetic (A) and bacterial killing (B) of naïve WT and PCV13-immunized WT or µMT mice during the first 30 min post i.v. infection with 10^6^ CFU of TIGR4. n = 3. (**C-F**) Abundance-dependent activation of pathogen capture in the liver by purified IgG. IgG antibodies were purified from natural or PCV13 immune serum and detected of IgG band by SDS-PAGE (C). 10^6^ CFU of TIGR4 were pre-incubated with different amounts of purified IgG before i.v. inoculated into naïve mice to test bacteremia kinetic (D), organ bacteria (E) and bacterial killing (F) during the first 30 min post infection. n = 3. (**G-I**) Vaccine abundance-dependent activation of endothelial capture of pneumococci. Mice were immunized once with various doses of PCV13 before being tested for the production of anti-CPS4 IgG as in Fig. 4A (G), bacterial capture by ECs and KCs as in Fig. 3F (H), and survival post i.p. infection with 10^6^ CFU of TIGR4 (I). Bacterial capture is demonstrated in **Movie S5**. n = 3-6 for (G) and 5-6 for (I); quantitation in (H) was shown as pooled data of 5-13 FOVs from two mice in each group. (**J-L**) Importance of boost immunization in PCV13-activated hepatic immunity. Mice were immunized once, twice or three times with 2.5 µl PCV13 before being tested for the production of anti-CPS4 IgG (J), bacterial capture by ECs and KCs as in Fig. 3F (K), and survival post i.p. infection with 10^7^ CFU of TIGR4 (L). Bacterial capture is demonstrated in **Movie S5**. n = 2-14 for (J) and 5 for (L); quantitation in (K) was shown as pooled data of 10-24 FOVs from two mice in each group. Data shown in (G and J) were representative of duplicate experiments; dotted line indicates the limit of detection; data are presented as the mean ± SEM. Statistics: (A and D) two-way ANOVA; (B, F, G and J) unpaired two-tailed Student’s t test; (I and L) log-rank (Mantel-Cox) test.

**Fig. S5.**
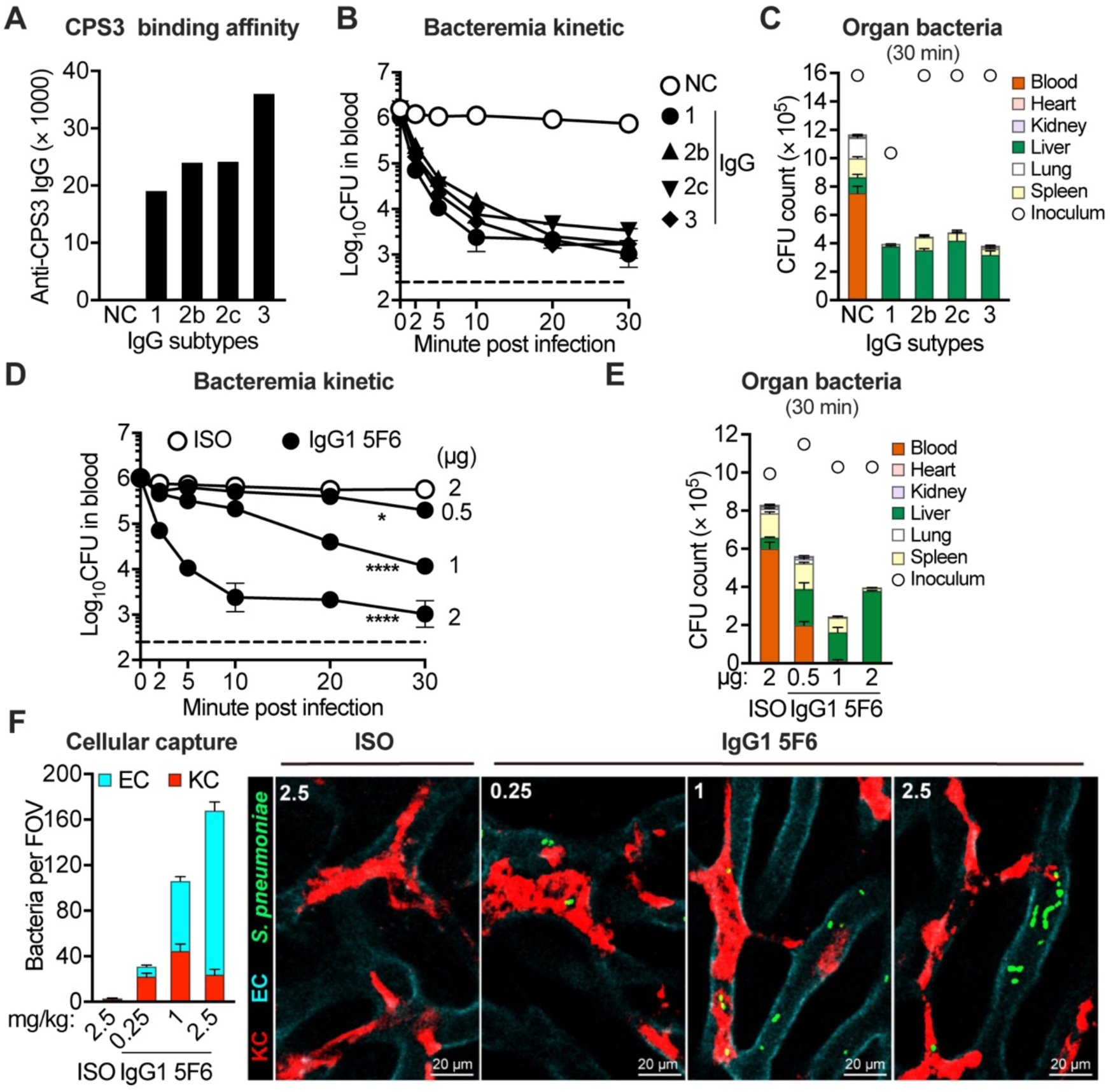
The antibody subtypes-dependent endothelial capture of pneumococci. (**A**) Anti-CPS3 IgG titer detection of recombinant monoclonal antibody variants with different IgG subtypes. The initial concentration was 0.5 mg/ml of purified antibody. (**B** and **C**) Bacteremia kinetic (B) and organ bacteria (C) in naïve mice post i.v. inoculated with 10^6^ CFU of serotype-3 pneumococci ST865 which were pretreated with 2 µg of different anti-CPS3 IgG antibody variants. n = 3. (**D-F**) IgG1 antibody abundance-dependent activation of endothelial capture of pneumococci. Mice were i.v. inoculated with 10^6^ CFU of ST865, which were pretreated with different amounts of anti-CPS3 IgG1 5F6 (0.5, 1 and 2 μg) or isotype control (ISO, 2 μg) to assess the bacteremia kinetic (D) and organ bacteria (E). Mice were i.v. inoculated with 0.25, 1 and 2.5 mg/kg of anti-CPS3 IgG1 5F6 immediately after i.v. injection of 5 × 10^7^ CFU of ST865 to assess bacterial capture by ECs and KCs as in Fig. 3F (F). Bacterial capture is demonstrated in **Movie S6**. n = 3 for (D and E); quantitation in (F) was shown as pooled data of 10 FOVs from two mice in each group. Data shown in (A) were performed once; dotted line indicates the limit of detection; data are presented as the mean ± SEM. Statistics: (D) two-way ANOVA.

**Fig. S6.**
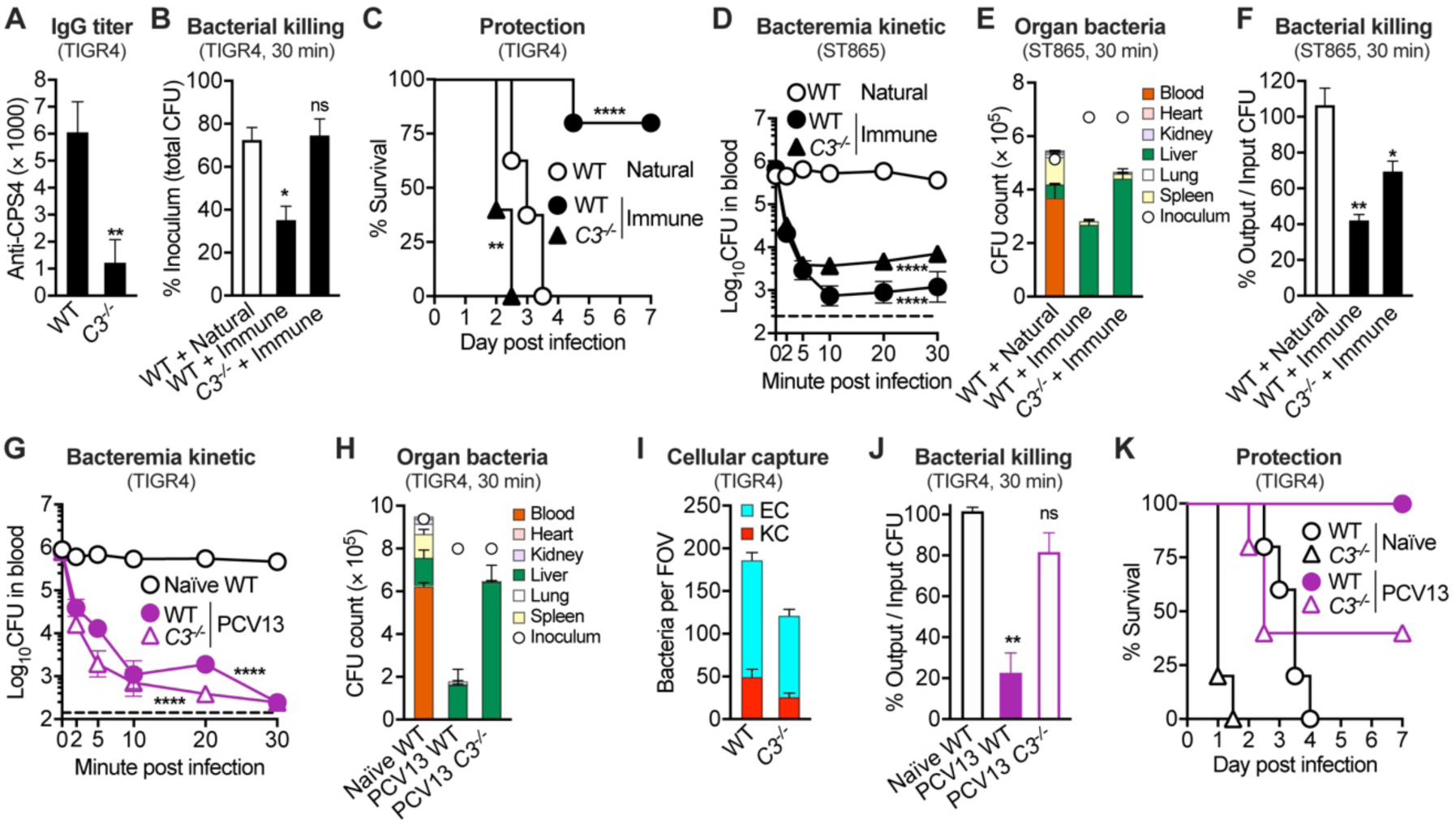
Role of complement system for PCV13-elicited pathogen clearance. (**A**) Anti-CPS4 IgG titer in sera of PCV13-immunized WT and *C3*^-/-^ mice. (**B** and **C**) Importance of complement C3 in PCV13 immune serum-mediated killing of ECs-captured TIGR4. Bacterial killing (B) and survival (C) in WT and *C3*^-/-^ mice after i.v. infection with natural or PCV13 immune serum-opsonized TIGR4 as in Fig. 4F. n = 3 for (B) and 5-10 for (C). (**D-F**) Dispensability of complement C3 for hepatic capture but essential for killing of ST865. Bacteremia kinetic (D), organ bacteria (E) and bacterial killing (F) of WT or *C3*^-/-^ mice at 30 min after i.v. infection of 10^6^ CFU of natural or PCV13 immune serum-opsonized ST865 as in Fig. 4F. n = 3. (**G**-**I**) Dispensability of complement C3 in PCV13 vaccine-activated capture of TIGR4 in the liver. Bacteremia kinetic (G), organ bacteria (H) and bacterial capture by ECs and KCs (I) of naïve WT and PCV13-immunized WT or *C3*^-/-^ mice post i.v. infection with of TIGR4 were presented as in Fig. 4F-H. Bacterial capture is demonstrated in **Movie S7**. n = 4-6 for (G and H); quantitation in (I) was shown as pooled data of 10-21 FOVs from two mice in each group. (**J** and **K**) Importance of complement C3 in PCV13 vaccine-activated killing of TIGR4. Bacterial killing (J) and survival (K) in naïve WT and PCV13-immunized WT or *C3*^-/-^ mice after i.v. infection with 10^6^ CFU of TIGR4. n =3 for (J) and 5 for (K). Data shown in (A) were representative of duplicate experiments; dotted line indicates the limit of detection; data are presented as the mean ± SEM. Statistics: (A, B, F and J) unpaired two-tailed Student’s t test; (C) log-rank (Mantel-Cox) test; (D and G) two-way ANOVA.

**Fig. S7.**
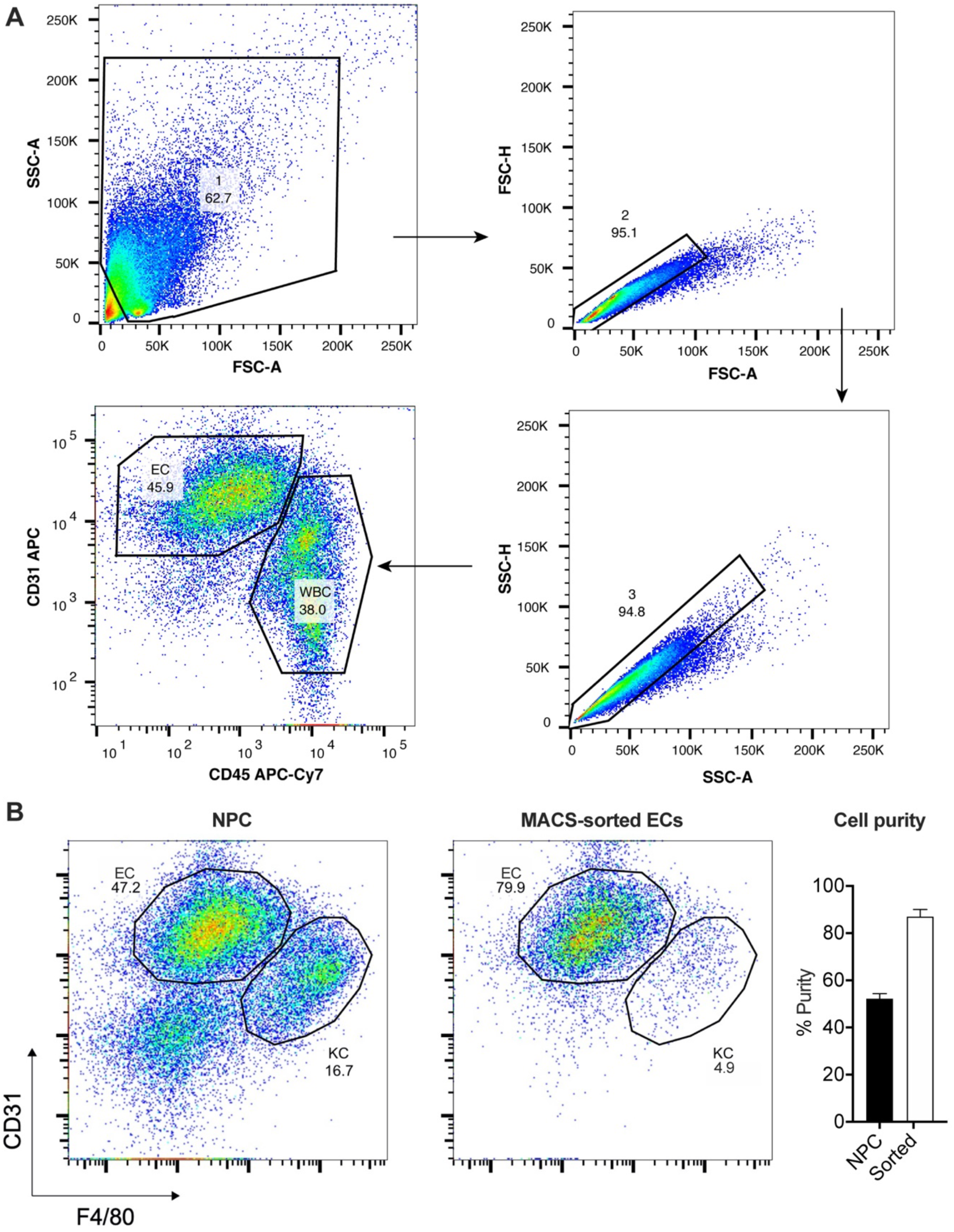
Gating strategy and cell purity detection of isolated ECs by flow cytometry. (**A**) For flow cytometry analysis, murine liver non-parenchymal cells were gated according to the forward- and side-scatter channel values. Doublets were excluded based on the FSC-A/H and SSC-A/H plotting. Liver ECs were identified as CD31^+^CD45^-^. n = 3. (**B**) ECs were isolated from murine liver non-parenchymal cells by magnetic-activated cell sorting (MACS) as described in Methods, and represented by CD31^+^CD45^-^ populations in the pre- or post-isolation. The cell purity was calculated by dividing positively labeled cells by total cells (left panel). Data shown above were representative of duplicate experiments; data are presented as the mean ± SEM.

**Fig. S8.**
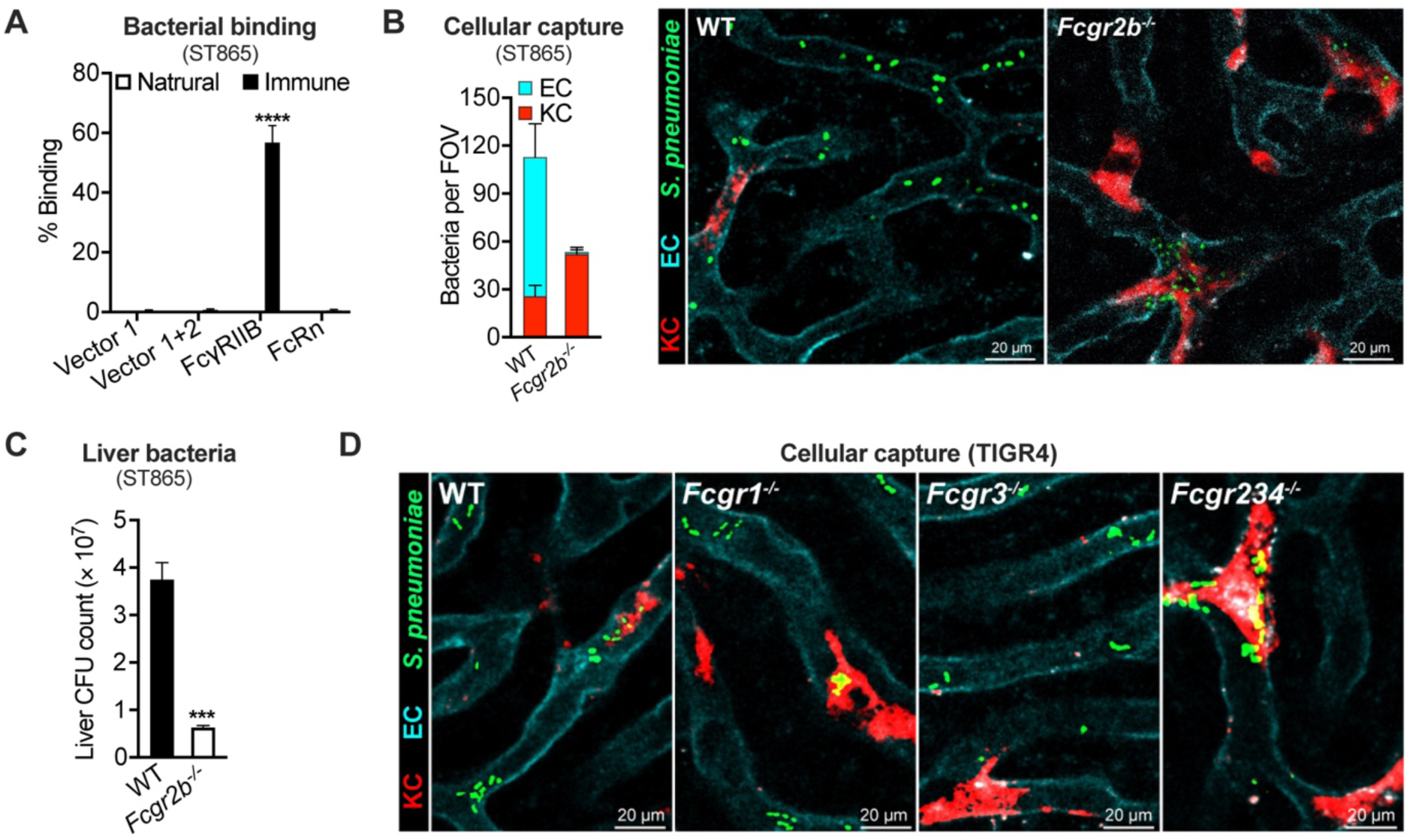
Importance of FcγRIIB in endothelial capture of antibody-opsonized bacteria. (**A**) *In vitro* binding of ST865 to CHO cells expressing FcγRIIB (*Fcgr2b*) or FcRn (*Fcgrt* and *B2m*) as in Fig. 5C. n = 3. (**B**) ECs- and KCs-associated ST865 in PCV13-immunized WT or *Fcgr2b*^-/-^ mice are presented as in Fig. 5F. Bacterial capture is demonstrated in **Movie S3**. Quantitation was shown as pooled data of 5 FOVs from two mice in each group. (**C**) Significant impairment of IgG1-mediated pathogen captures in the livers of *Fcgr2b*^-/-^ mice. WT and *Fcgr2b*^-/-^ mice were i.v. infected with 5 × 10^7^ CFU of ST865 before inoculation of 2.5 mg/kg of anti-CPS3 IgG1 5F6; bacteria recovered from the liver at 30 min post infection was presented. n = 3. (**D**) Contribution of FcγRs importance in PCV13-elicited pathogen capture by KCs. Representative IVM images of the liver sinusoids are shown to demonstrate that multiple FcγRs (especially FcγRIII) are required for the KC’s capture of pneumococci (TIGR4) in PCV13-immunized WT, *Fcgr1^-/-^*, *Fcgr2b^-/-^*, *Fcgr3^-/-^* and *Fcgr2b/3/4^-/-^*mice. Related data are also presented elsewhere in Fig. 5M. Data shown in (A) were representative of duplicate experiments; data are presented as the mean ± SEM. Statistics: (A) two-way ANOVA; (C) unpaired two-tailed Student’s t test.

**Fig. S9.**
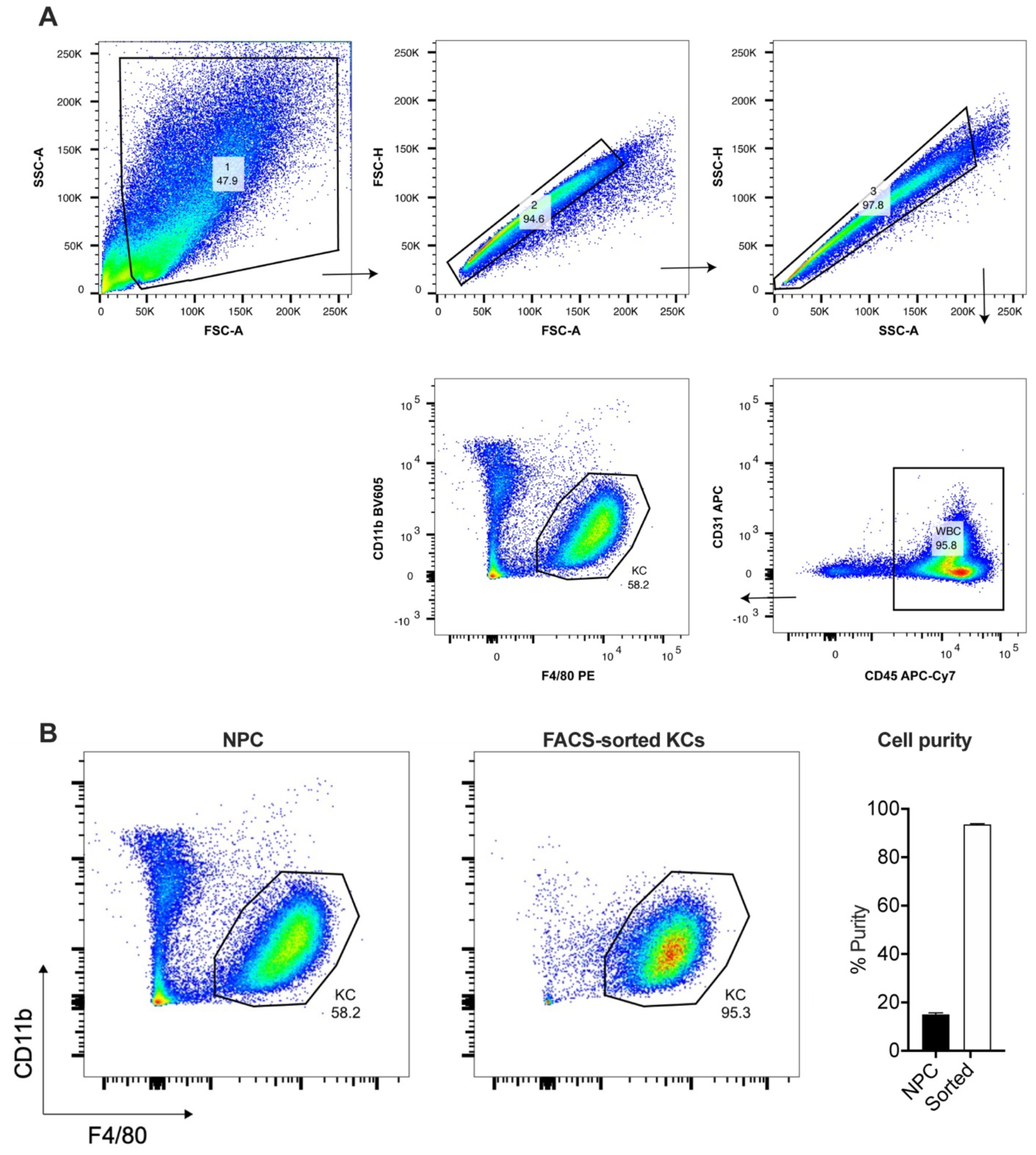
Gating strategy and cell purity detection of isolated KCs by flow cytometry. (**A**) For flow cytometry analysis, murine liver non-parenchymal cells were gated according to the forward- and side-scatter channel values. Doublets were excluded based on the FSC-A/H and SSC-A/H plotting. Liver KCs were identified as CD45^+^CD31^-^CD11b^low^F4/80^+^. n = 3. (**B**) KCs were isolated from murine liver non-parenchymal cells by fluorescence activated cell sorting (FACS) as described in Methods, and represented by CD45^+^ CD11b^low^ F4/80^+^ populations in the pre- or post-isolation. The cell purity was calculated by dividing positively labeled cells by total cells (left panel). Data shown above were representative of duplicate experiments; data are presented as the mean ± SEM.

**Fig. S10.**
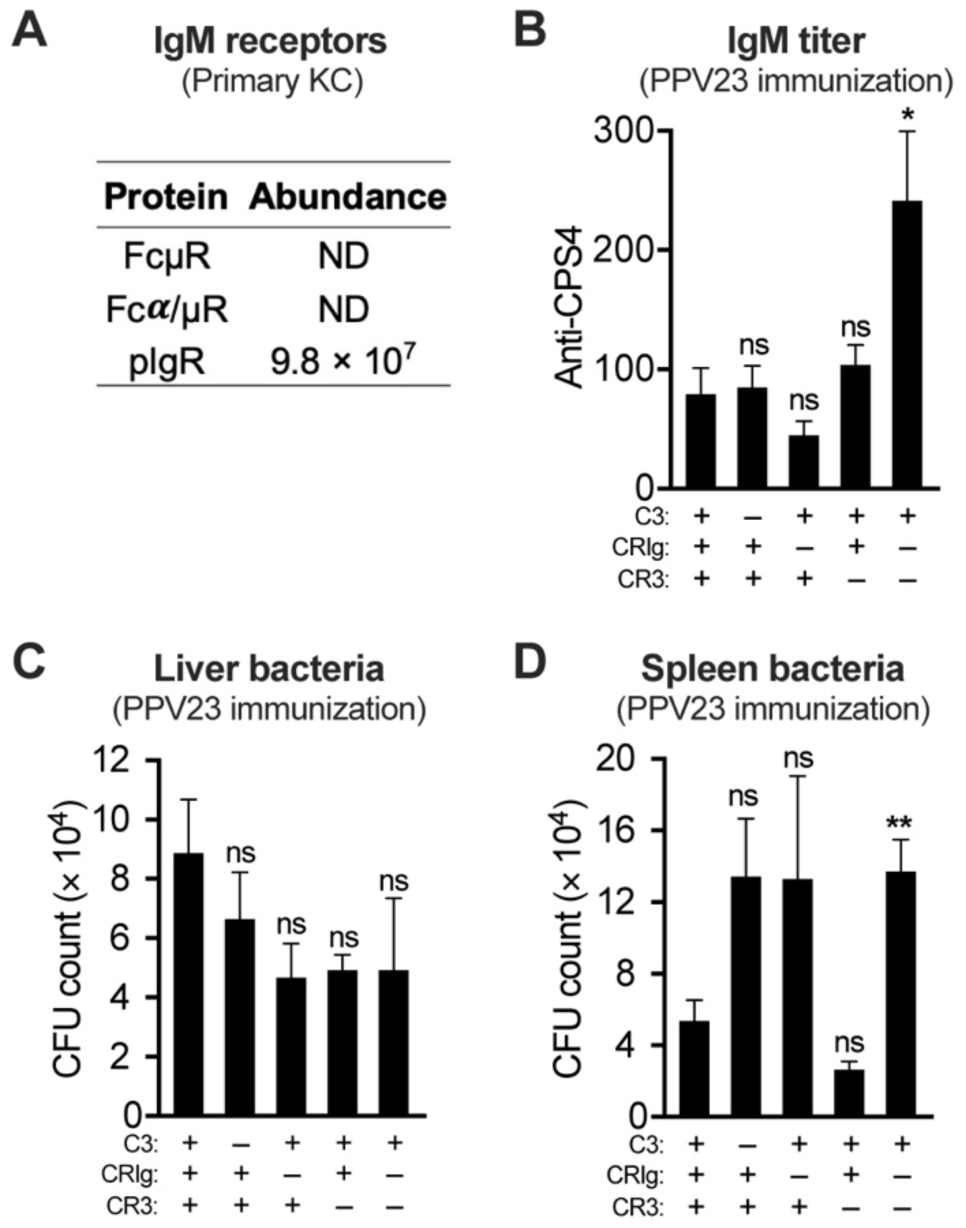
Role of complement system for PPV23-elicited pathogen clearance by Kupffer cells. (**A**) Relative abundance of each IgM receptors in primary murine KCs as detected by mass spectrometry as in Fig. 5J. Additional data are available in **Table S2**. (**B**-**D**) Importance of complement system in PPV23-activated pathogen clearance. Anti-CPS4 IgM titer was detected in sera of PPV23-immunized WT, C3- and C3 receptors- (CRIg and CR3) deficient C57BL/6J mice by ELISA (B); n = 7-12. The roles of complement C3 and C3 receptors (CRIg and CR3) in pathogen capture by liver (C) and spleen (D) in PPV23-immunized WT (n = 4), *C3^-/-^* (n = 5), CRIg KO (n = 6), CR3 KO (n = 4) and CRIg/CR3 KO (n = 5) C57BL/6J mice were shown as the CFU count in blood. Data shown in (A) were representative of duplicate experiments; data are presented as the mean ± SEM. Statistics: (B-D) unpaired two-tailed Student’s t test.

**Fig. S11.**
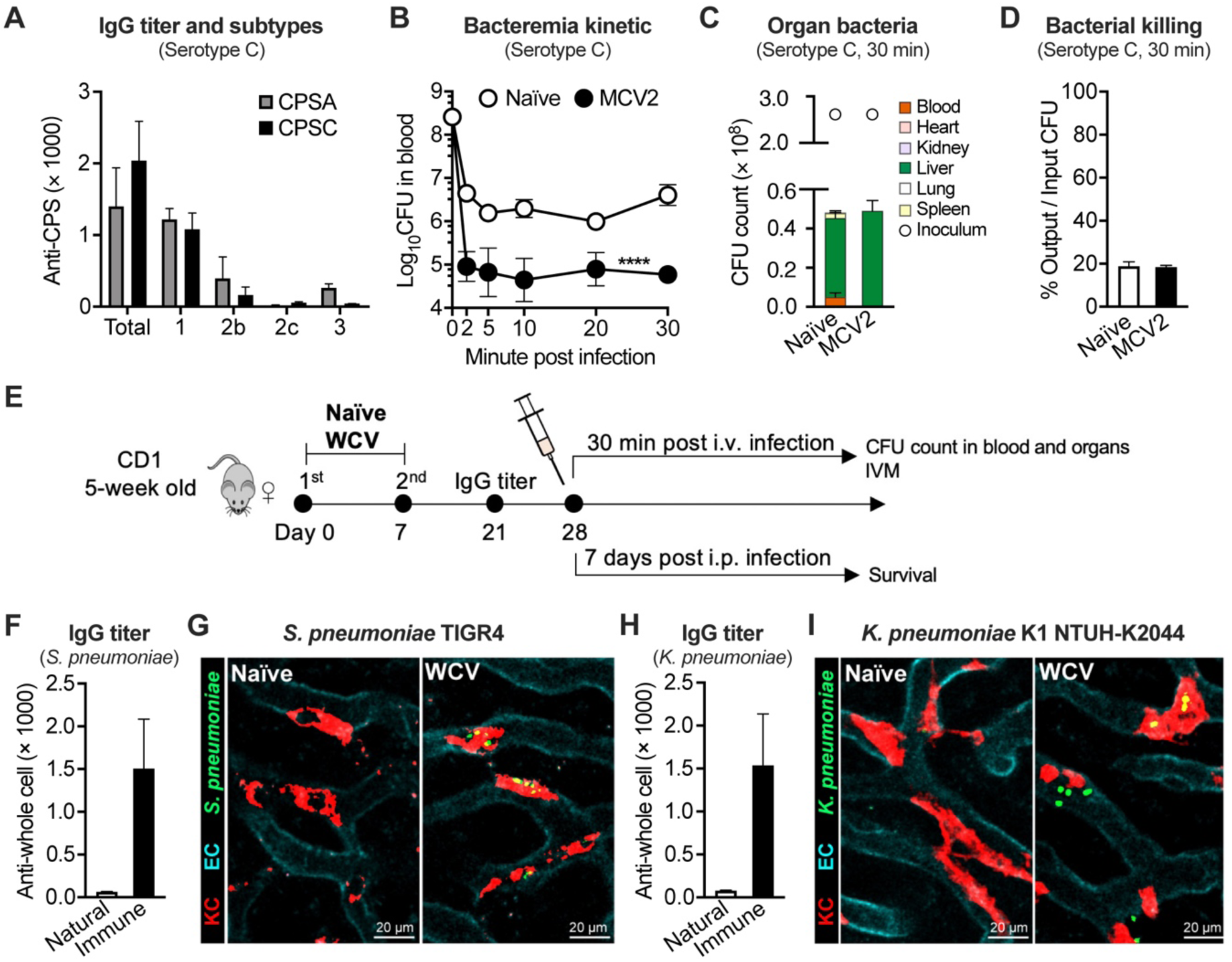
Vaccine-induced pathogen capture by liver for other virulent human pathogens. (**A**) Anti-CPS IgG titer and subtypes in the sera of *Neisseria meningitidis* conjugate vaccine MCV2-immunized (immune) and naïve (natural) mice. n = 2-3. (**B** to **D**) Bacteremia kinetic (B), organ bacteria (C) and bacterial killing (D) of naïve and MCV2-immunized mice after i.v. infection with 2 × 10^8^ CFU of serotype-C *N. meningitidis*. n = 3. (**E**) Experimental design. 5-week-old female CD1 mice were subcutaneously immunized with 10^8^ CFU of inactivated whole-cell bacteria (WCV) or remained untreated (naïve) on day 0, followed by once consecutive boosts on day 7. All animals were tested for serum antibodies response on day 21 before i.v. challenged to test bacteremia kinetic and organ bacteria and cellular capture by ECs and KCs during the first 30 min post infection; i.p. challenged with three doses of bacteria to monitor the survival for 7 days. (**F** and **G**) Whole cell vaccine induced immunoprotection of *S. pneumoniae*. Mice were immunized with 10^8^ CFU of inactivated TIGR4 to assess the production of anti-whole cell IgG (F) and bacterial capture by ECs and KCs as in Fig. 3L (G), Bacterial capture is demonstrated in **Movie S11**. (**H** and **I**) Whole cell vaccine induced immunoprotection of *K. pneumoniae*. The anti-whole cell IgG (H) and bacterial capture by ECs and KCs (I) were assessed as in (F and G) expect for using WCV and pathogen of *K. pneumoniae*. Bacterial capture is demonstrated in **Movie S11**. Data shown in (A) were representative of duplicate experiments; data are presented as the mean ± SEM. Statistics: (B) two-way ANOVA.

### Supplementary Tables S1-S4

**Table S1.** The mass spectrometry data of membrane proteins isolated from murine ECs.

**Table S2.** The mass spectrometry data of membrane proteins isolated from murine KCs.

**Table S3.** Mice strains and antibodies used in this study.

**Table S4.** Bacterial strains, plasmids and primers used in this study.

### Supplementary Movies S1 to S11

**Movie S1. Vaccine-activated capture of serotype-4 pneumococci in liver sinusoids.** Liver sinusoid endothelial cells (ECs, cyan) and resident macrophage Kupffer cells (KCs, red) capture of i.v. inoculated serotype-4 pneumococci (5 × 10^7^ CFU, strain TIGR4, green) from the bloodstream of naïve, PCV13- and PPV23-immunized mice. Intravital microscopic visualization was initiated immediately before bacterial injection; imaging data at the onset of bacterial appearance in the circulation are presented. The playback rate is eight frames per second. The quantitative data were shown in Fig. 3F and 3K; selected images in Fig. 3L.

**Movie S2. The role of major phagocytes for vaccine-activated capture of serotype-4 pneumococci in liver sinusoids**. Visualization of EC- and KC-associated pneumococci in the liver sinusoids of naïve, PCV13- and PPV23-immunized mice were imaged as in Movie S1 without or with phagocytes depletion by CLL/Gr1 treatment. The quantitative data were shown in Fig. 3F and 3K; selected images in Fig. 3L.

**Movie S3. Vaccine-activated capture of serotype-3 pneumococci in liver sinusoids**. Visualization of EC- and KC-associated pneumococci in the liver sinusoids were imaged as in Movie S1 of naïve, PCV13- and PPV23-immunized mice without or with CLL/Gr1 treatment, and PCV13-immunized WT or *Fcgr2b*^-/-^ mice after i.v. inoculation of 5 × 10^7^ of serotype-3 pneumococci ST865. The quantitative data and selected images are shown in fig. S3F, S3J, S3K and S8B.

**Movie S4**. **Dose-dependent activation of endothelial capture of serotype-4 pneumococci by PCV13 immune serum.** Visualization of ECs- and KCs-associated pneumococci in the liver sinusoids of naïve mice were imaged as in Movie S1 after i.v. inoculation of 2 × 10^7^ CFU of serum-opsonized TIGR4. The bacteria were pre-incubated with 100, 200 and 300 µl of heat-inactivated PCV13 immune serum or 300 µl of heat-inactivated natural serum for 30 min before injection. The quantitative data and selected images are shown in Fig. 4C.

**Movie S5**. **Vaccine dose-dependent activation of endothelial pathogen capture**. Visualization of ECs- and KCs-associated pneumococci in the liver sinusoids of mice vaccinated once with various doses of PCV13 or vaccinated once, twice or three times with 2.5 µl PCV13 were imaged as in Movie S1 after i.v. inoculation of 5 × 10^7^ of TIGR4. Selected images are shown in fig. S4I and S4L.

**Movie S6**. **Abundance- and subtypes-dependent impact of serotype-specific monoclonal antibody on endothelial pathogen capture**. Naïve mice were i.v. inoculated with different amounts of anti-CPS3 IgG1 5F6 antibody or anti-CPS3 IgG antibodies with different IgG subtypes (2.5 mg/kg) immediately after i.v. injection of 5 × 10^7^ CFU of ST865 to assess pathogen capture by ECs and KCs as in Movie S1. IVM imaging was also presented in *Fcgr2b*^-/-^ mice with 2.5 mg/kg of anti-CPS3 IgG1 5F6 IgG1 injection after infection as in Movie S1. The quantitative data and selected images are shown in Fig. 4E, 5G, 5I and fig. S5F.

**Movie S7. No requirement of complement for PCV13-activated capture of serotype-4 pneumococci in liver sinusoids**. Visualization of EC- and KC-associated pneumococci in the liver sinusoids of WT or *C3*^-/-^ mice were imaged as in Movie S4 after i.v. inoculated with serum-opsonized of TIGR4. The bacteria were pre-treated with 300 µl of heat-inactivated PCV13 immune for 30 min before injection. IVM imaging was also presented in PCV13-immunized WT or *C3*^-/-^ mice as in Movie S1 after i.v. inoculated with TIGR4. The quantitative data and selected images are shown in fig. 4H and S6I.

**Movie S8**. **Importance of FcγRIIB in vaccine-elicited capture of serotype-4 pneumococci in liver sinusoids.** Visualization of EC- and KC-associated pneumococci in the liver sinusoids of PCV13-immunized WT, *Fcgr2b*^-/-^ or *Fcgrt*^-/-^ mice were imaged as in Movie S1 after i.v. inoculation of 5 × 10^7^ of TIGR4. The quantitative data are shown in Fig. 5F; Selected images in Fig. 5H.

**Movie S9**. **The requirement of complement and complement receptor (CRIg and CR3) for PPV23-activated capture of serotype-4 pneumococci by Kupffer cell in liver sinusoids**. Visualization of ECs- and KCs-associated pneumococci in the liver sinusoids of PPV23-immunized WT, *C3*^-/-^, CRIg KO, CR3 KO and CRIg/CR3 KO mice were imaged as in Movie S1 after i.v. inoculation of 5 × 10^7^ of TIGR4. The quantitative data were shown in Fig. 6E; selected images in Fig. 6F.

**Movie S10. Importance of FcγRIIB in vaccine-elicited capture of serotype-C meningococci in liver sinusoids**. Visualization of EC- and KC-associated meningococci in the liver sinusoids of naïve WT and MCV2-immunized WT or *Fcgr2b*^-/-^ mice were imaged as in Movie S1 after i.v. inoculation of 5 × 10^7^ of serotype-C meningococci. Selected images are shown in Fig. 7A.

**Movie S11**. **Whole cell vaccine induced capture of *S. pneumoniae* and *K. pneumoniae* by KCs in liver sinusoids**. Visualization of ECs- and KCs-associated bacteria in the liver sinusoids of naïve and serotype-4 *S. pneumoniae* or serotype-K1 *K. pneumoniae* NTUH-K2044 whole cell vaccine (WCV)-immunized mice were imaged as in Movie S1 after i.v. inoculation of 5 × 10^7^ of *S. pneumoniae* or *K. pneumoniae*, respectively. The quantitative data were shown in Fig. 7E and 7I; Selected images in fig. S11G and S11I, respectively.

