## Supplementary material for "Potent Bacterial Vaccines Require FcγRIIB-mediated Pathogen Capture by Liver Sinusoidal Endothelium": Table S3

**Table S3. Mice strains and antibodies used in this study**

| **STRAIN** | **SOURCE** | **IDENTIFIER** |
| --- | --- | --- |
| C57BL/6 µMT KO | Jackson Laboratory | RRID: IMSR_JAX:002288 |
| C57BL/6 *C3^-/-^* | Jackson Laboratory | RRID: IMSR_JAX:003641 |
| C57BL/6 CRIg KO | (*1*) | RRID: MGI:3806097 |
| C57BL/6 *Fcgr1^-/-^* | Model Organisms Center | RRID: IMSR_NM-KO-18033 |
| C57BL/6 *Fcgr2b^-/-^* | Jackson Laboratory | RRID: IMSR_JAX:002848 |
| C57BL/6 *Fcgr3^-/-^* | Model Organisms Center | RRID: IMSR_NM-KO-190189 |
| C57BL/6 *Fcgr2b/3/4^-/-^* | Model Organisms Center | RRID: IMSR_NM-KO-18037 |
| C57BL/6 *Fcgrt^-/-^* | Model Organisms Center | RRID: IMSR_NM-KO-00133 |
| C57BL/6 CR3 KO | This study | N/A |
| C57BL/6 CRIg/CR3 KO | This study | N/A |
| **ANTIBODY** | **SOURCE** | **IDENTIFIER** |
| APC-Cy7 anti-mouse CD45 (Clone: 30-F11) | BioLegend | RRID: AB_312981 |
| APC anti-mouse CD31 (Clone: 390) | BioLegend | RRID: AB_312904 |
| PE anti-mouse F4/80 (Clone: BM8) | BioLegend | RRID: AB_893486 |
| BV605 anti-mouse CD11b (Clone: M1/70) | BioLegend | RRID: AB_2565431 |
| FITC anti-mouse Ly6G (Clone: 1A8) | BD Pharmingen | RRID: AB_394207 |
| eF450 anti-mouse Ly6C (Clone: HK1.4) | Invitrogen | RRID: AB_10805519 |
| AF647 anti-mouse F4/80 (Clone: BM8) | BioLegend | RRID: AB_893492 |
| AF594 anti-mouse CD31 (Clone: MEC13.3) | BioLegend | RRID: AB_2563319 |
| PE anti-mouse Ly6G (Clone: 1A8) | BioLegend | RRID: AB_1186099 |
| Anti-mouse Ly6G/Ly6C (Clone: NIMP-R14) | BioXCell | RRID: AB_2819047 |
| Rat IgG2b isotype control (Clone: LTF-2) | BioXCell | RRID: AB_1107780 |
| Horseradish peroxidase (HRP)-conjugated goat anti-mouse IgG | EasyBio | Cat# BE0102-100 |
| Horseradish peroxidase (HRP)-conjugated goat anti-mouse IgM | Elabscience | Cat# E-AB-1008 |
| Horseradish peroxidase (HRP)-conjugated goat anti-mouse IgG1 | Proteintech | RRID: AB_2890964 |
| Horseradish peroxidase (HRP)-conjugated goat anti-mouse IgG2b | Proteintech | RRID: AB_2890966 |
| Horseradish peroxidase (HRP)-conjugated goat anti-mouse IgG2c | Proteintech | RRID: AB_2890967 |
| Horseradish peroxidase (HRP)-conjugated goat anti-mouse IgG3 | Proteintech | RRID: AB_2890968 |
| Mouse IgG1 isotype control (Clone: MOPC-21) | BioXcell | RRID: AB_1107784 |
| Mouse anti-His tag (Clone: OTI2B5) | ZSGB-BIO | Cat# TA-02 |
| Anti-β-actin antibody | Beyotime | Cat# AF0003 |
