## Supplementary material for "Potent Bacterial Vaccines Require FcγRIIB-mediated Pathogen Capture by Liver Sinusoidal Endothelium": Table S4

**Table S4. Bacterial strains, plasmids and primers used in this study**

| **Bacterial strains** | | | | |
| --- | --- | --- | --- | --- |
| Strain ID | Species | Serotype | Source | Region |
| TH865 | *Streptococcus pneumoniae* | 3 | Cerebrospinal Fluid | USA |
| TIGR4 |  | 4 | Blood | Norway |
| TH2884 |  | 8 | Blood | China |
| TH16187 | *Neisseria meningitidis* | C | Cerebrospinal Fluid | China |
| TH12813 | *Staphylococcus aureus* | - | Blood | China |
| NTUH-K2044 | *Klebsiella pneumoniae* | K1 | Liver abscesses | China |

| **Plasmids** | | | | | |
| --- | --- | --- | --- | --- | --- |
| Plasmid ID | Backbone | Insertion segments | Primers for  segments | Template  DNA | Digestion |
| pTH14790 | pCDH | cDNA of murine *fcgr2b* with C-terminal 6 x His tag | Pr16958/Pr16959 | cDNA library of EC | XbaI/EcoRI |
| pTH14792 | pCDH | cDNA of murine *fcgrt* with C-terminal 6 x His tag | Pr16962/Pr16963 |  | XbaI/EcoRI |
| pTH14793 | pFUGW | cDNA of murine *belat2mG* with C-terminal 6 x His tag | Pr16964/Pr16965 |  | XbaI/NheI |
| pTH15735 | pCDH | cDNA of murine *fcgr1* with C-terminal 6 x His tag | Pr18178/Pr18179 | cDNA library of KC | XbaI/EcoRI |
| pTH14791 | pCDH | cDNA of murine *fcgr3* with C-terminal 6 x His tag | Pr16960/Pr16961 |  | XbaI/EcoRI |
| pTH15736 | pCDH | cDNA of murine *fcgr4* with C-terminal 6 x His tag | Pr18180/Pr18181 |  | XbaI/EcoRI |
| pTH14794 | pFUGW | cDNA of murine *fcgγ* with C-terminal 6 x His tag | Pr16966/Pr16967 |  | XbaI/NheI |
| pTH16703 | H vector: 5F6 IgG1 | Heavy chain of variable (VH) and constant (CH_IgG1_) regions of 5F6 IgG1 | Pr19037/Pr19038 (VH) | GenBank: DQ228147.1 | AgeI/NotI |
|  |  |  | Pr19049/Pr19050 (CH_IgG1_) | GenBank: MH208238.1 |  |
|  |  |  | Pr19037/Pr19050 (Fusion) | VH+ CH_IgG1_ |  |
| pTH16668 | H vector: 5F6 IgG2b | Heavy chain of variable (VH) and constant (CH_IgG2b_) regions of 5F6 IgG2b | Pr19037/Pr19038 (VH) | GenBank: DQ228147.1 | AgeI/NotI |
|  |  |  | Pr19039/Pr19040 (CH_IgG2b_) | GenBank: LC052323.1 |  |
|  |  |  | Pr19037/Pr19040 (Fusion) | VH+ CH_IgG2b_ |  |
| pTH16669 | H vector: 5F6 IgG2c | Heavy chain of variable (VH) and constant (CH_IgG2c_) regions of 5F6 IgG2c | Pr19037/Pr19038 (VH) | GenBank: DQ228147.1 | AgeI/NotI |
|  |  |  | Pr19041/Pr19042 (CH_IgG2c_) | GenBank: LC037230.1 |  |
|  |  |  | Pr19037/Pr19042 (Fusion) | VH+ CH_IgG2c_ |  |
| pTH16670 | H vector: 5F6 IgG3 | Heavy chain of variable (VH) and constant (CH_IgG3_) regions of 5F6 IgG2c | Pr19037/Pr19038 (VH) | GenBank: DQ228147.1 | AgeI/NotI |
|  |  |  | Pr19043/Pr19044 (CH_IgG3_) | GenBank: MT019336.1 |  |
|  |  |  | Pr19037/Pr19044 (Fusion) | VH+ CH_IgG3_ |  |
| pTH16671 | L vector: 5F6 | Light chain of variable (VL) and constant (CL) regions of 5F6 | Pr19045/Pr19046 (VL) | GenBank: DQ228148.1 | AgeI/NotI |
|  |  |  | Pr19047/Pr19048 (CL) | GenBank: MH208239.1 |  |
|  |  |  | Pr19045/Pr19048 (Fusion) | VL+ CL |  |

| **Primers** | |
| --- | --- |
| Primer ID | Sequence (5’-3’) |
| Pr16958 | GCTCTAGAATGGGAATCCTGCCGTTCCTACTGA |
| Pr16959 | CGGAATTCCTAATGGTGATGGTGATGATGAATGTGGTTCTGGTAATCATGCTCT |
| Pr16960 | GCTCTAGAATGTTTCAGAATGCACACTCTGGAA |
| Pr16961 | CGGAATTCTCAATGGTGATGGTGATGATGCTTGTCTTGAGGAGCCTGGTGCTTT |
| Pr16962 | GCTCTAGAATGGGGATGCCACTGCCCTG |
| Pr16963 | CGGAATTCTCAATGGTGATGGTGATGATGGGAAGTGGCTGGAAAGGCATTTGCA |
| Pr16964 | GCTCTAGAATGGCTCGCTCGGTGACCCT |
| Pr16965 | GAGAGCTAGCTCAATGGTGATGGTGATGATGCATGTCTCGATCCCAGTAGACGGTC |
| Pr16966 | GCTCTAGAATGATCTCAGCCGTGATCTTGTTCT |
| Pr16967 | GAGAGCTAGCCTAATGGTGATGGTGATGATGCTGGGGTGGTTTCTCATGCTTCAGA |
| Pr18178 | GCTCTAGAATGATTCTTACCAGCTTTGGAGATG |
| Pr18179 | CGGAATTCTCAATGGTGATGGTGATGATGACTTTGGGAAGTTTGTGCCCCAGTA |
| Pr18180 | GCTCTAGAATGTGGCAGCTACTACTACCAACAG |
| Pr18181 | CGGAATTCTCAATGGTGATGGTGATGATGCTTGTCCTGAGGTTCCTTGCTCCAT |
| Pr19037 | TTTTTCTAGTAGCAACTGCAACCGGTGTACATTCTCAGGTCCAACTGCAGCAG |
| Pr19038 | GGCCCTTGGTGGTGGCGCTCGAGGCTGCAGAGACAGTGACCAG |
| Pr19039 | TCACTGTCTCTGCAGCCTCGAGCGCTAAAACAACACCCCCATCA |
| Pr19040 | GCCAAGCTTGGGAGCGGCCGCTCATTTACCCGGAGACCGG |
| Pr19041 | TCACTGTCTCTGCAGCCTCGAGCGCCAAAACAACAGCCCCAT |
| Pr19042 | CCAAGCTTGGGAGCGGCCGCTCATTTACCCAGAGACCGGG |
| Pr19043 | GTCACTGTCTCTGCAGCCTCGAGCGCTACAACAACAGCCCCA |
| Pr19044 | GCCAAGCTTGGGAGCGGCCGCTCATTTACCAGGGGAGCG |
| Pr19045 | TTTTCTAGTAGCAACTGCAACCGGTGTACATTCTGATGTTGTGGTGACCCAAAC |
| Pr19046 | TACAGTTGGTGCAGCATCCGTACGCGTACGCTTGATTTCCAGC |
| Pr19047 | CTGGAAATCAAGCGTACGCGTACGGATGCTGCACCAACTGTAT |
| Pr19048 | GCGGCCAAGCTTGGGAGCGGCCGCTCAACACTCATTCCTGTTG |
| Pr19049 | CTGGTCACTGTCTCTGCAGCCTCGAGCGCCACCACCAAGGGCCCATCT |
| Pr19050 | CATGGCGGCCAAGCTTGGGAGCGGCCGCTCATTTACCAGGAGAGTGGG |
